# MiR-302-Induced anti-aging neural stem cells enhance cognitive function and extend lifespan

**DOI:** 10.1101/2023.02.13.528232

**Authors:** Yuanyuan Li, Jing Sun, Yuanyuan Zheng, Tingting Xu, Yanan Zhang, Yuesi Wang

## Abstract

Neural stem cells play a vital role in maintaining tissue stability and extending lifespan. Transplanting these cells to treat neurodegenerative diseases faces challenges like cellular aging, low viability, and immune rejection. We have effectively reprogrammed human fibroblasts into induced neural stem cells (iNSCs) via a single-factor miR-302a strategy, which converted skin fibroblasts into human-induced neural stem cells (hiNSCs) within 2-3 days. These cells showed delayed aging and increased resistance to oxidative stress compared to wild-type cells. Implanting them into the hippocampus of senescence-accelerated mice improved cognitive performance in severe Alzheimer’s, prolonged lifespan by 34%, increased fatigue resistance, and improved hair regeneration and reproductive capacity. Our findings suggest that miR-302a-hiNSCs can improve functional recovery in Alzheimer’s and promote healthy aging.

## INTRODUCTION

The process of aging is a multifaceted and intricate phenomenon that is characterized by a chronic dysregulation of cellular behavior, leading to a decline in tissue and organ function, which culminates in a plethora of degenerative diseases, including Alzheimer’s disease, debility, and tumors (*1, 2*). Despite the decline in adult neurogenesis, which is the process of generating neurons in the adult brain, it continues to persist throughout life and is a critical component in the occurrence of diseases such as Alzheimer’s syndrome. As a result, it serves as a potential target for prolonging cognitive health and reducing the risk of neurodegenerative diseases. With the increasing life expectancy, the prevalence of these health challenges is becoming more apparent, emphasizing the need for effective interventions (*3–6*). The age-related loss of neural stem cell numbers and/or activity is believed to play a pivotal role in the decline of brain function (*5, 7–10*). Hippocampal neural stem cells (NSCs) are particularly susceptible to cellular senescence (*11, 12*). and disruption of the persistence of these cells in the hypothalamus has been shown to accelerate aging in mice (*13*). Therefore, interventions aimed at reversing neural stem cell aging may increase the period of cognitive health in humans.

Some researchers have suggested that a decline in the function of the central nervous system (CNS) is one of the principal markers of aging in mammals (*14*). Although there has been considerable advancement in the field and continuous study, a cure for this chronic illness still has not been discovered. As a result, such declines in cell number or function may be susceptible to interventions aimed at slowing or preventing aging while also delaying or preventing other diseases and thus extending a healthy lifespan. An attractive theory is to say that the decline in brain function with age is the result of loss of stem cell numbers and/or activity, then an intervention that supplements stem cell numbers or reverses stem cell aging may extend the human lifespan and health span (*15*).

Stem cell transplantation appears to be a promising therapeutic option for neurodegenerative disorders such as AD and aging because it has the potential to exert multiple restorative effects, including cell replacement and paracrine effects (*16, 17*). However, most preclinical studies and clinical trials use allogeneic mesenchymal stem cells (MSCs), NSCs, or induced pluripotent stem cells (iPSCs), autologous patient-derived neural stem cell therapies may have significant advantages for clinical use. The ability of autologous neural stem cell therapy to avoid immune rejection not only eliminates the complications of immunosuppressive therapy, but also prolongs the persistence of NSCs to increase the visibility of the cells and the durability of the therapy. Research advances in direct cellular reprogramming offer the possibility of developing patient-specific cellular therapies. Induced neural stem cells (iNSCs) have been derived by reprogramming somatic cells with multiple transcription factors and other methods to avoid the pluripotent phase of induced pluripotent stem cells and the risks associated with low differentiation efficiency and carcinogenicity (*18–26*). Jing Nai-Ho et al used PB MNC cells reprogrammed as induced neural stem cells to rescue AD cognitive deficits by strengthening synaptic networks. However, However, the impact of iNSCs treatment generated by direct reprogramming on the health span of older organisms has not been explored (*27*).

As a first step towards the development of personalized human-induced neural stem cells (hiNSCs) therapies, we show that a single miR-302a reprogramming strategy can efficiently and quickly convert human skin fibroblasts into iNSCs in as little as 2-3 days, as evidenced by surface marker, molecular, and functional analysis of the cells. We then show that miR-302a-iNSCs have better anti-aging and antioxidant functions than other neural stem cell sources. Here, we show that miR-302a-hiNSC hippocampal transplantation significantly improves cognitive function in rapidly aging older SAMP8 mice. Elderly mice receiving miR-302a-hiNSC showed lower levels of frailty and significant improvements in lifespan extension, running wheel testing, glucose regulation, fur regeneration, and renal function. In addition, SAMP8 mice transplanted with iNSCs showed some improvement in hearing compared to the control SAMP8 group. In this study, we sought to identify strategies based on miR-302a reprogramming to obtain iNSCs with high antioxidant and anti-aging capacity to provide a faster and more efficient method of iNSCs acquisition for application in research and clinical therapy.

## RESULTS

### Single microRNAs rapidly and efficiently convert human fibroblasts into iNSCs

The current study aimed to explore the potential of a single miR-302a component of the miR-302/367 cluster in reprogramming human skin fibroblasts into induced neural stem cells (iNSCs). Previous literature has demonstrated the advantageous effects of combining the miR-302/367 cluster in improving the efficiency of reprogramming techniques (*28, 29*). To determine if the overexpression of a single component of the miR-302 cluster can reprogram fibroblasts, we generated lentiviral/adenoviral vectors that encoded the miR-302a sequence, a member of the miR-302 cluster and used it to reprogram human foreskin-derived fibroblasts (HFFs) obtained from postnatal sources. Surprisingly, we discovered that simply overexpressing miR-302a converted human skin fibroblasts into NSCs quickly and efficiently. HFFs (26- and 29-year-old males) and eyelid fibroblasts (41-year-old female) were isolated and identified as Vimentin positive and negative on Nestin, GFAP, and TUJ1 by immunostaining or RT-qPCR (Data not shown).

The HFFs, eyelid fibroblasts, and mouse cell line L929 cells were then infected with either a lentiviral or an adenovirus vector expressing miR-302a (Fig. 1A). For one day, we cultured HFFs on gelatin-coated coverslips. Within 24 to 48 hours of miR-302a overexpression in HFFs, their morphology changes rapidly, and clusters form. The earliest appearance of a colony was observed at 13 hours (Fig. 1B). Three days after transfection with miR-302a, the colony formation rate of HFFs reached 1.11%, then increased to 2.29% on the fifth day and 8.02% on the seventh day. (Fig. 1C). Eyelid fibroblasts, like HFFs, can form neurospheres rapidly and have a higher colony-forming rate (Data not shown), demonstrating that this approach to generating neurospheres from human fibroblasts is quicker and more effective than any currently existing method (Table S1).

**Fig. 1.**
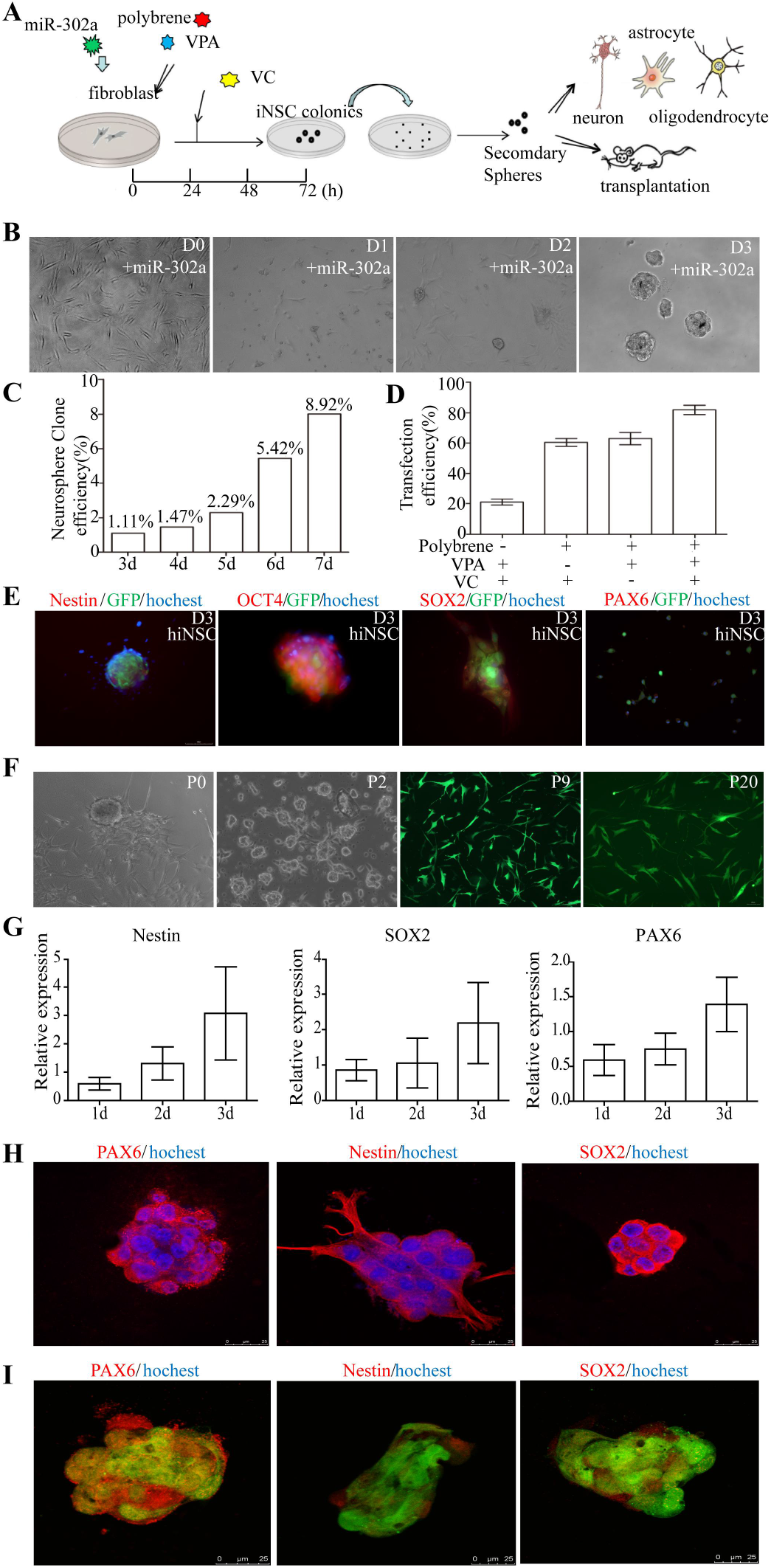
Generation and characterization of iNSCs from human postnatal foreskin-derived fibroblasts (HFFs). (**A**) Schematic representation of the reprogramming process with single microRNAs miR-302a to generate neurospheres from human fibroblasts. (**B**) Phase-contrast images of HFFs after overnight treatment with miR-302a virus vector in fibroblast medium. Has-miR-302a-infected cells spontaneously formed neurosphere-like colonies on gelatin-coated glass coverslip from 1-3 days after transfection. (**C**) The colony formation efficiency of neurospheres generated by hiNSCs from human fibroblasts at 3, 4, 5, 6, and 7 days after miR-302a transfection. (**D**) Effect of the three small molecules used to increase transfection efficiency for reprogramming with has-miR-302a. Data are presented as mean ± SD (n=3). (**E**) Expression of Nestin, OCT4, SOX2, and PAX6 in hiNSCs neurospheres at day 3 after miR-302a transfection. (**F**) Neurospheres derived from miR-302a reprogrammed human fibroblasts can be expanded from P0 to P20 passages with adherent and suspension culture alternately. (**G**) RT-qPCR reveals that hiNSCs expressed typical NSC markers Nestin, SOX2, and PAX6 on day 3. (**H**) The control HFFs infected with Ptf1a lentiviruses derived neurospheres were PAX6, Nestin, and SOX2 positive expression in Immunostaining. (**I**) Neurospheres induced from HFFs by miR-302a were immunoreactive for PAX6, Nestin, and SOX2.

Within three days, almost all of the transfected cells had generated neurospheres. The reprogramming efficiency can be improved by adding varying VPA, VC, and polybrene concentrations. Using GFP-tagged transfected cells, the optimal combination of the three concentrations showed a virus-cell transfection efficiency of up to 90%. (Fig. 1D).

Zhang et al identified a cocktail of nine components that could efficiently reprogram identified mouse fibroblasts into a neural stem cell-like cells using the percentage of SOX2+/Nestin+ positive cells as a criterion for reprogramming efficiency (*30*). INSCs expressed Nestin and SOX2 markers in immunofluorescence and RT-qPCR results (Fig. 1, G and I). Based on the findings, we concluded that miR-302a transfected fibroblasts had a 90% reprogramming efficiency for neural stem cells. The transfection method with mir-302a for reprogramming fibroblasts into NSCs is faster and more effective than any other existing reprogramming method. (Table S1).

In the colonies, the expression of crucial NSC markers, Nestin, SOX2, and PAX6, was detectable within 72 hours, as shown by immunostaining results, and increased gradually over time with prolonged culture. (Fig. 1E). On day 3, 90% of surviving infected fibroblasts were found to be Nestin positive by flow cytometry. These fibroblasts were reprogrammed into NSCs with a higher level of purity and quantity (Data not shown).

The induced neurospheres were harvested, broken down, and re-cultured in an NSC expansion medium for suspension and monolayer culture. Starting on day 2, the new iNSCs secondary neurospheres quickly developed. To enlarge and purify neural stem cells, suspension culture and monolayer culture are alternately used. For more than 10 months and more than 20 passes, the iNSCs could be expanded and maintained steadily. The morphologies of the iNSCs in passages 1, 2, 9, and 20 were uniform and stable (Fig. 1F), and they were uniformly positive for Nestin, SOX2, and PAX6 (Fig. 1G), demonstrating their capacity for self-renewal.

RT-qPCR analysis showed that the expression of Nestin, PAX6, and SOX2 in HFFs undergoing reprogramming into NSCs started low on the first day and increased progressively over time (Fig. 1G). hiNSCs reprogrammed on glass slides expressed Nestin, PAX6, and SOX2 more strongly than cells connected directly to culture plates, and neurosphere development occurred more quickly. This outcome is quite comparable to that published by Bag óJR et al in 2017 (*31*). OCT4 is not expressed during hiNSCs differentiation, hence the hiNSCs did not have a high OCT4 expression.

The study also found that single miR-302a molecules can efficiently transform mouse L929 cells into NSCs, with neurospheres visible after just three days and around 80% of cells having round-sized and clustered morphologies (Fig. 4A). The expression levels of Nestin, PAX6, and SOX2 in the reprogrammed cells were comparable to those in the SC029 control mouse NSC lines. Furthermore, this method proved faster and more efficient than existing approaches(Fig. 4, B to E). Overall, these results show that single miR-302a molecules can efficiently and quickly transform huge numbers of fibroblasts from mice and humans into neural stem cells (Table S2).

**Fig. 2.**
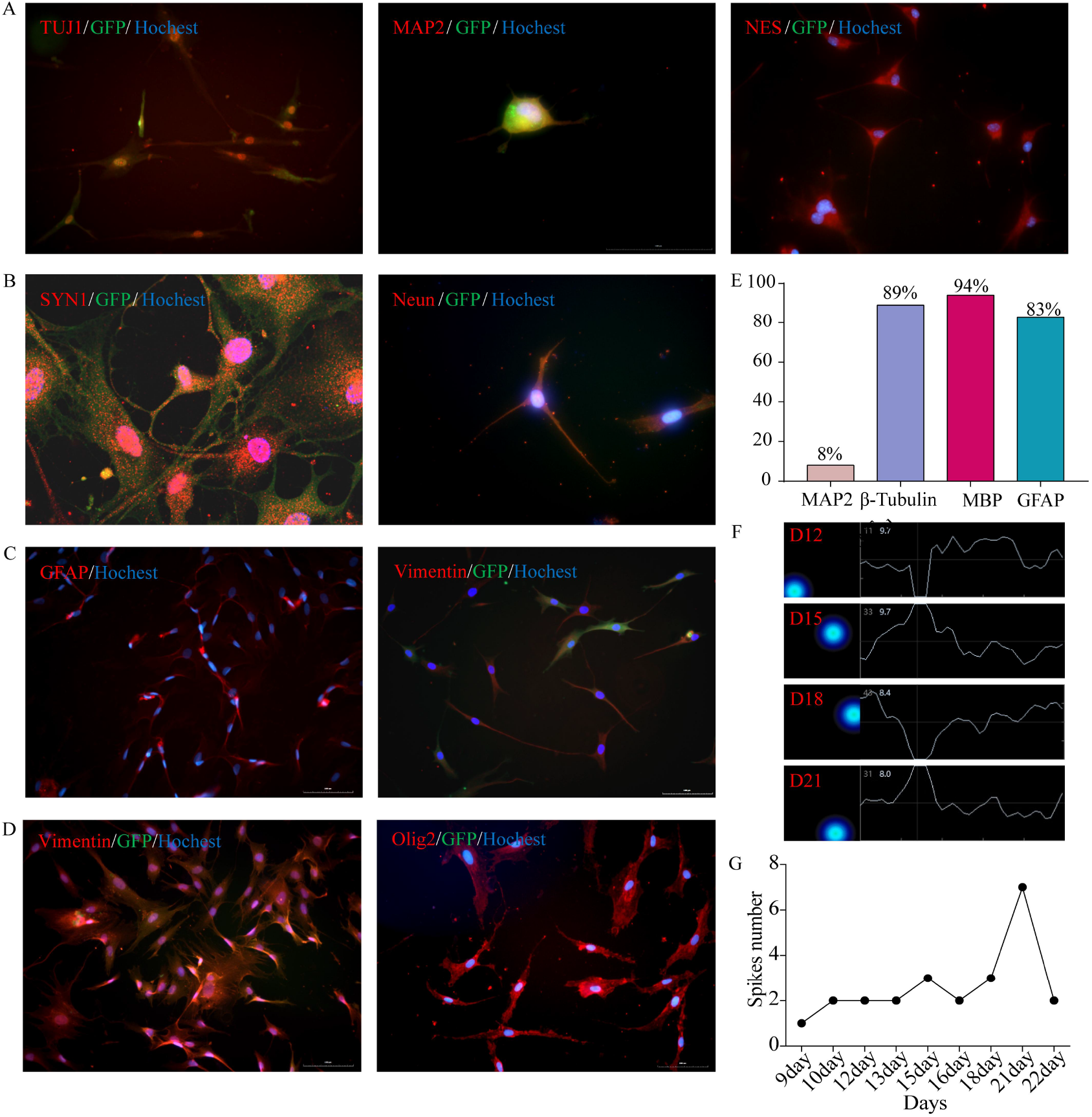
Multipotency of iNSCs in vitro. (**A**) hiNSCs could be differentiated into TUJ1+, MAP2+ and NES+ neurons by day 8 in culture after growth factor withdrawal. (**B**) hiNSCs were able to differentiate into SYN1+ and NeuN+ neurons. (**C**) hiNSCs differentiate into subtypes of astrocytes, including GFAP+ and Vimentin+ positive cells. (**D**) hiNSCs can robustly generate MBP+ and Olig2+ oligodendrocytes by day 22 in vitro. Scale bars represent 100 μM in A-D. (**E**) Quantification of MAP2, β-Tubulin, MBP and GFAP positive cells rate differentiated from hiNSCs. (**F**) Cumulative activity map and spiking amplitude changes of differentiated neurons derived from hiNSCs at D12, D15, D18 and D21 using the MEA detection. (**G**) Quantification of the number of spike-positive electrodes of differentiated neurons with spontaneous action potential derived from hiNSCs on MEA increased from D12 to D21, and dropped on D22.

**Fig. 3.**
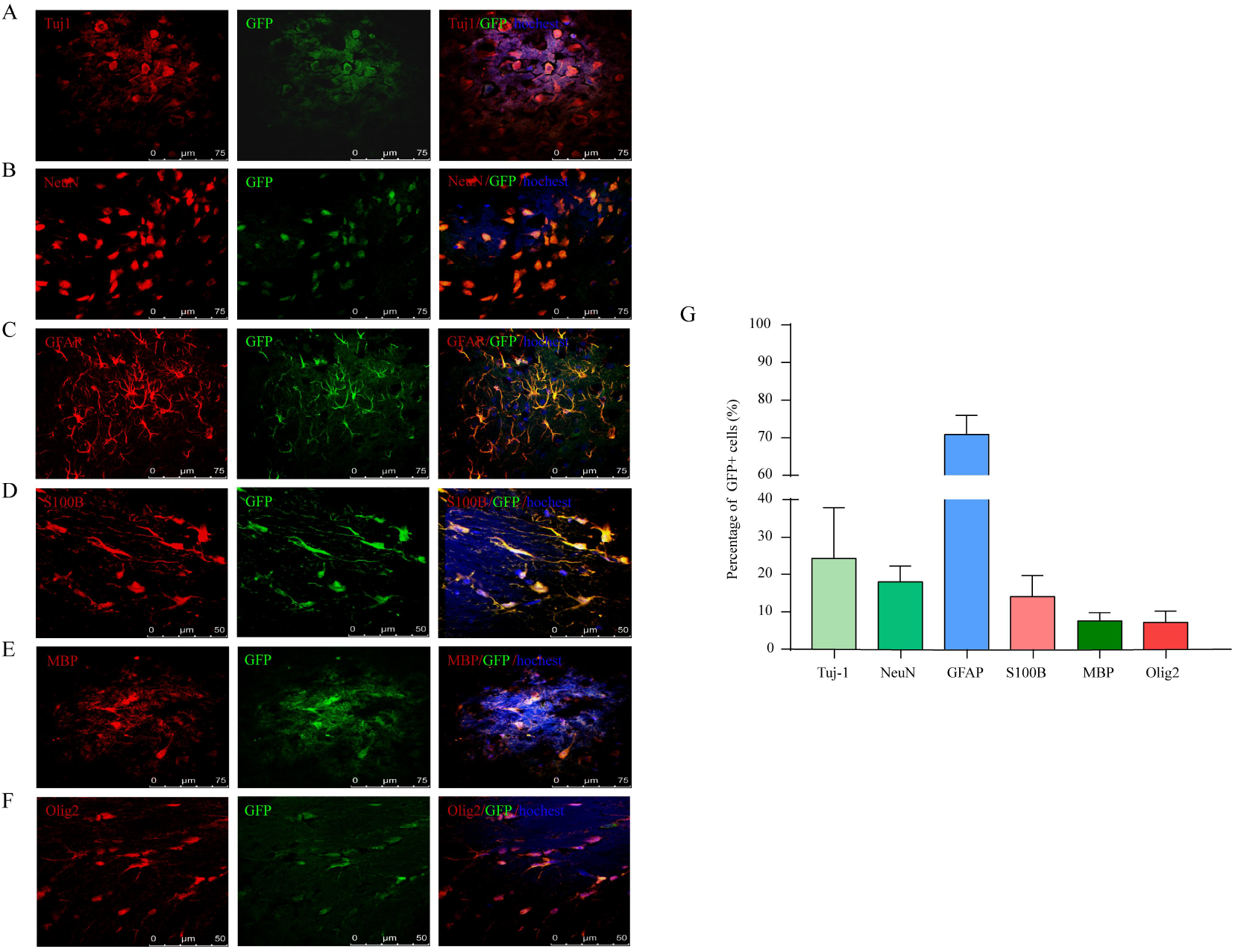
Mulitipotency of miR-302a-reprogrammed iNSCs in vivo. The miR-302a-reprogrammed iNSCs were injected into the striatum of 2-month-old nude mice (n=8). 3 weeks post-transplantation, GFP+ iNSCs migrated and integrated into the mice brain. (**A** and **B**) Immunostains reveal that hiNSCs can differentiate into TUJ1+ and NeuN+ neurons. (**C** and **D**) GFP+ iNSC can differentiate into GFAP+ and S100b+ astrocytes. (**E** and **F**) Injected GFP+ cells co-expressing the oligodendrocytes cell marker MBP and Olig2. (**G**) Quantification of labeled cells in the sections showed that GFP+ cells differentiated into TUJ1+, NeuN+, GFAP+, S100b+, MBP+, and Olig2+ cells. Scale bars represent 75 μM in (A), (B), (C), and (E), and 50 μM in (D) and (F).

**Fig. 4.**
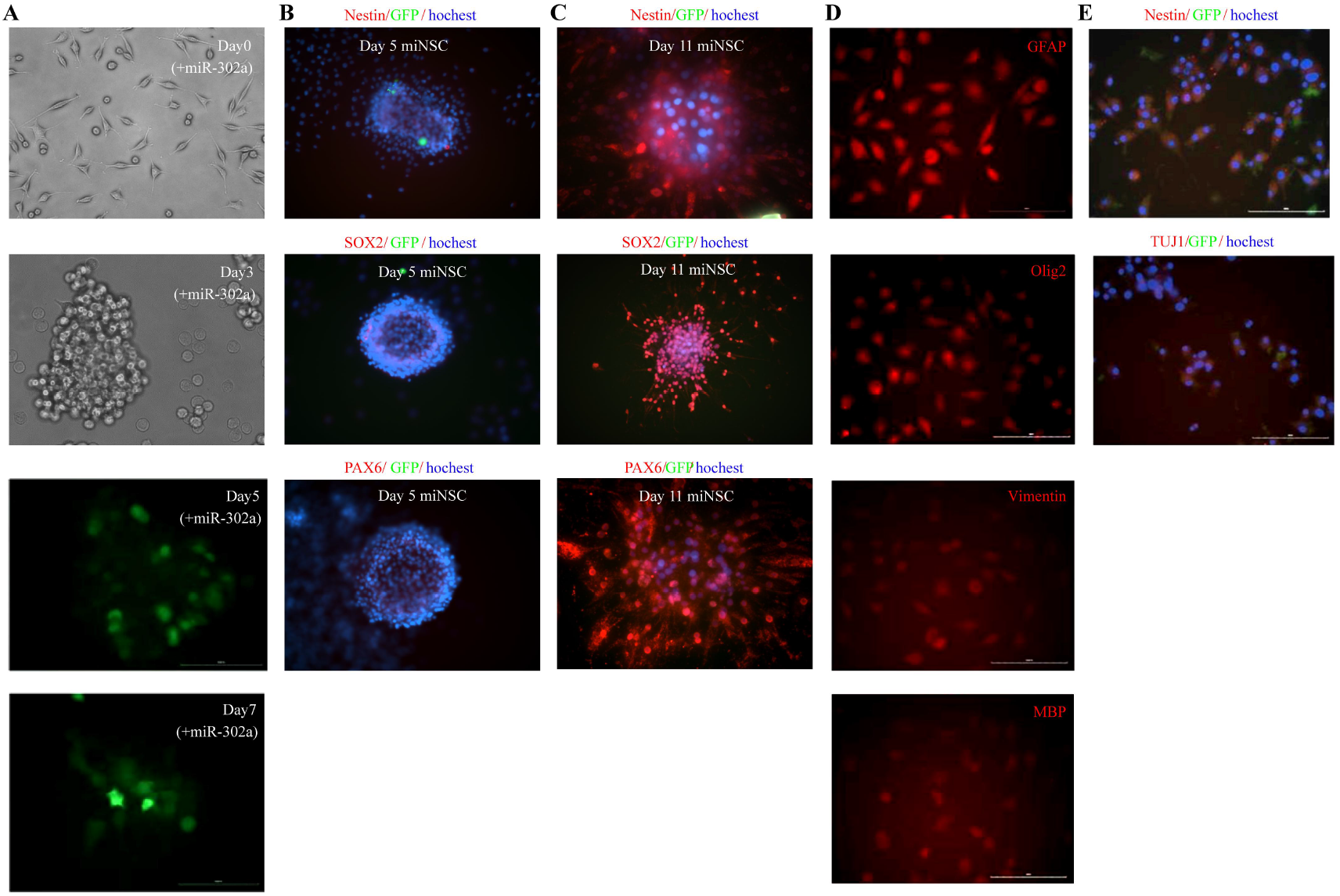
MiR-302a reprograms mouse L929 fibroblasts into miNSCs. (**A**) Representative fields show a time course of colony formation at days 0, 3, 5, and 7 after mouse L929 single-cell plating with miR-302a transfection. (**B**) Immunofluorescent staining of mouse L929 fibroblasts-miNSCs for Nestin, SOX2, and PAX6 on day 5. (**C**) Immunofluorescence staining for Nestin, SOX2, and PAX6 of miNSCs induced by mouse L929 fibroblasts produced on day 11 d after secondary spheroid formation in culture. (**D**) Representative images of in vitro differentiation of miNSCs into astrocytes (GFAP+ and Vimentin+) and oligodendrocytes (MBP+ and Oligo2+). (**E**) Representative images of in vitro differentiation of miNSCs into MAP2+ and TUJ1+ neurons. Data shown are representative images from three independent experiments which gave similar results.

### Multipotency of hiNSCs in vitro

The ability of iNSCs to produce oligodendrocytes, astrocytes, and neurons served as a measure of their developmental potential. iNSCs were grown on feeders for three days to induce spontaneous differentiation. After removing the feeders, bFGF, and EGF, the iNSCs differentiated into a substantial number of TUJ1-positive neurons within four days. iNSCs differentiated into neurons that were positive for TUJ1, Map2, Nestin, SYN1, and NeuN after a further one to two weeks of culture (Fig. 2, A and B). A quantitative analysis conducted after three weeks showed that 89% of the cells were positive for TUJ1. (Fig. 2E). GFAP and Vimentin were both presents in the iNSC-derived astrocytes, according to immunocytochemistry (Fig. 2C). Additionally, during the differentiation process into oligodendrocytes, the hiNSCs were also able to differentiate into cells that were positive for MBP and Olig2. (Fig. 2D). 92% of the cells underwent astrocyte differentiation and became GFAP-positive astrocytes. 94% of developed cells in the oligodendrocyte differentiation media were MBP-positive (Fig. 2E). In iNSCs from various sources of fibroblasts, the expression patterns of the critical surface markers are comparable. These findings firmly establish the multipotency and ability of miR-302a-reprogrammed-iNSCs to generate a variety of neuronal subtypes.

The microelectrode array (MEA) analysis, a technique that measures the electrical activity of a cell population, was used to assess the electrophysiological function of neurons derived from iNSCs over a period of 9-21 days. The machine began recording spontaneous action potentials on day 9. The number of spontaneous action potentials grew gradually, peaked on day 21, and then started to decline on day 22. Until cells die, this pattern will persist (Fig. 2F). The neurons derived from iNSCs display a distinct spontaneous firing pattern in a synchronous cluster, indicating that the neural network has matured. After stimulation, the differentiated cells exhibit even more pronounced and robust excitatory action potentials. (Fig. 2G).

Furthermore, cells expressing the synaptic protein synapsin can be observed during neuronal differentiation, indicating their ability to establish connections between neurons (Fig. 2B). This demonstrates synaptic connections between the differentiated adult neurons. As a result, hiNSCs reprogrammed with miR-302a has the potential to develop into fully mature and functional neurons.

### INSCs can survive, integrate and differentiate in mice brain in vivo

The cortex and striatum of 3 to 5-week-old nude mice were microinjected with hiNSCs (Fig. 3 and fig. S1) and miNSCs (Fig. S2) to evaluate the in vivo developmental and differentiation capacity. Immunostaining showed that GFP-positive hiNSCs and miNSCs could be found and moved into the brain tissue adjacent to the implantation site after 2 to 4 weeks of transplantation. The transplanted hiNSCs and miNSCs are capable of giving rise to TUJ1 and NeuN-positive neurons, GFAP and S100b-positive astrocytes, and MBP and Olig2-positive glial cells. (Fig. 3, Fig. S1 and Fig. S2). The percentage of tagged cells in each region that differentiated into TUJ1+, NeuN+, GFAP+, S100b+, MBP+, and Olig2+ cells was 24.18%, 17.88%, 70.85%, 14.00%, 7.55%, and 7.20%. (Fig. 3G). The transplantation of miR-302a-induced NSCs has successfully developed neurons, astrocytes, and oligodendrocytes within the mice’s brains without causing any teratoma formation. Additionally, we examined many iNSCs from various passages; hippocampal teratoma development was not observed. These results indicate that miR-302a-hiNSCs possess the ability to survive and develop within the adult brain in vivo and the capacity to evade teratoma formation.

### Reprogramming of mouse fibroblasts by miR-302a into iNSCs

Mouse L929 cells were converted into neural stem cells by miR-302a using a procedure similar to HPF reprogramming to confirm that it can not only reprogram human fibroblasts but also murine fibroblasts (Fig. 4A). 90% of the cells were spherical, clumped, and forming neurospheres by day 3. High levels of Nestin, SOX2, and PAX6 expression were seen in miR-302a-reprogrammed miNSCs from day 5 to day 11 (Fig. 4, B and C). The miNSCs expressed Nestin, SOX2, and PAX6 in a manner that was comparable to that of the SCR029 mouse cell line used as a control. The differentiation of miR-302a-reprogrammed mouse neural stem cells (miNSCs) into neurons, astrocytes, and oligodendrocytes was further studied both in vitro and in vivo.

The miNSCs were differentiated into MAP2 and TUJ1 positive neurons, GFAP and Vimentin positive astrocytes, and MBP and Olig2 positive oligodendrocytes in vitro and in vivo. (Fig. 4, E and D). Transplanted miNSCs were injected into adult mice’s striatums and differentiated into neurons with TUJ1 and NeuN immunoreactivity, astrocytes with GFAP and S100b immunoreactivity, or oligodendrocytes with Olig2 immunoreactivity (Fig. S2, A to E). Quantification of the labeled cells in the sections revealed that, of the GFP+ cells, 22.27%, 19.51%, 72.77%, 23.17%, and 9.12% differentiated into TUJ1+, NeuN+, GFAP+, S100b+, and Olig2+ cells derived from miNSCs (Fig. S2G). This technique for reprogramming mouse neural stem cells is quicker and more effective than currently used techniques, similar to the creation of human iNSCs (Table S2). These results demonstrate the effectiveness and efficiency of single miR-302a molecules in converting large quantities of human and mouse fibroblasts into neural stem cells.

### Anti-aging and antioxidant effects of culture supernatant from miR-302a-hiNSCs on senescent skin fibroblasts and neural stem cells

We investigated whether the supernatant of cultured miR-302a-hiNSC had anti-senescence and oxidative effects on HPFF. HPFF, SB431542-iPSC-iNSC, and miR-302a-hiNSC cultured with supernatant of HPFF, miR-302a-hiNSCs were treated with 300 mM hydrogen peroxide for 24 h. The supernatant of HPFF, SB-iPSC-iNSC, and miR-302a-hiNSC was treated with 300 mM hydrogen peroxide for 24 h. Compared to HPFF, SB-iPSC-iNSC Supernatant, miR-302a-hiNSC Supernatant consistently exhibited reduced senescence-associated β-galactosidase (β-gal) activity. Compared with H_2_O_2_+HPFF, H_2_O_2_+HPFF+miR-302a-hiNSCs Supernatant exhibited reduced β-gal activity (Fig. 5, A and B), reduced induced apoptotic cells, and increased cell proliferation (Fig. 5, C to F). Next, we verified whether miR-302a-hiNSC itself has anti-aging and oxidative effects. Compared with miR-302a-hiNSC, SB-iPSC-iNSC, and H_2_O_2_-induced SB-iPSC-iNSC, H_2_O_2_-induced miR-302a-hiNSC consistently exhibited reduced senescence-associated β-galactosidase (β-gal) activity, with senescent cells reaching only 8.33%, while H_2_O_2_-induced SB-iPSC-iNSC senescent cells reached 46.67% (Fig. 5, G and I). Compared with miR-302a-hiNSCs, H_2_O_2_-induced apoptotic cells of miR-302a-hiNSCs reached only 29.67% (Fig. 5, H and J). Foxo3 regulates physiological oxidative stress response, Sirt6 overexpression delays aging, Nrf2 is a well-known transcription factor that regulates antioxidant expression, and we continued to investigate whether the three longevity factors Nrf2, Sirt6, and Foxo3 are expressed in miR-302a-hiNSCs. We found that miR-302a-hiNSCs significantly increased the expression of Nrf2, Sirt6, and Foxo3 compared with control HPFF and Ptf1a-hiNSCs, and further expressed that miR-302a-hiNSCs could delay cellular senescence (Fig. 5K). MiR-302a-hiNSCs significantly improved the anti-aging and antioxidant capacity of fibroblasts. MiR-302a-hiNSCs significantly improved the expression of Nrf2, Sirt6 and Foxo3. Cells with significantly improved anti-aging and antioxidant capacity compared to supernatants cultured from iPSC-iNSCs sources.

**Fig. 5.**
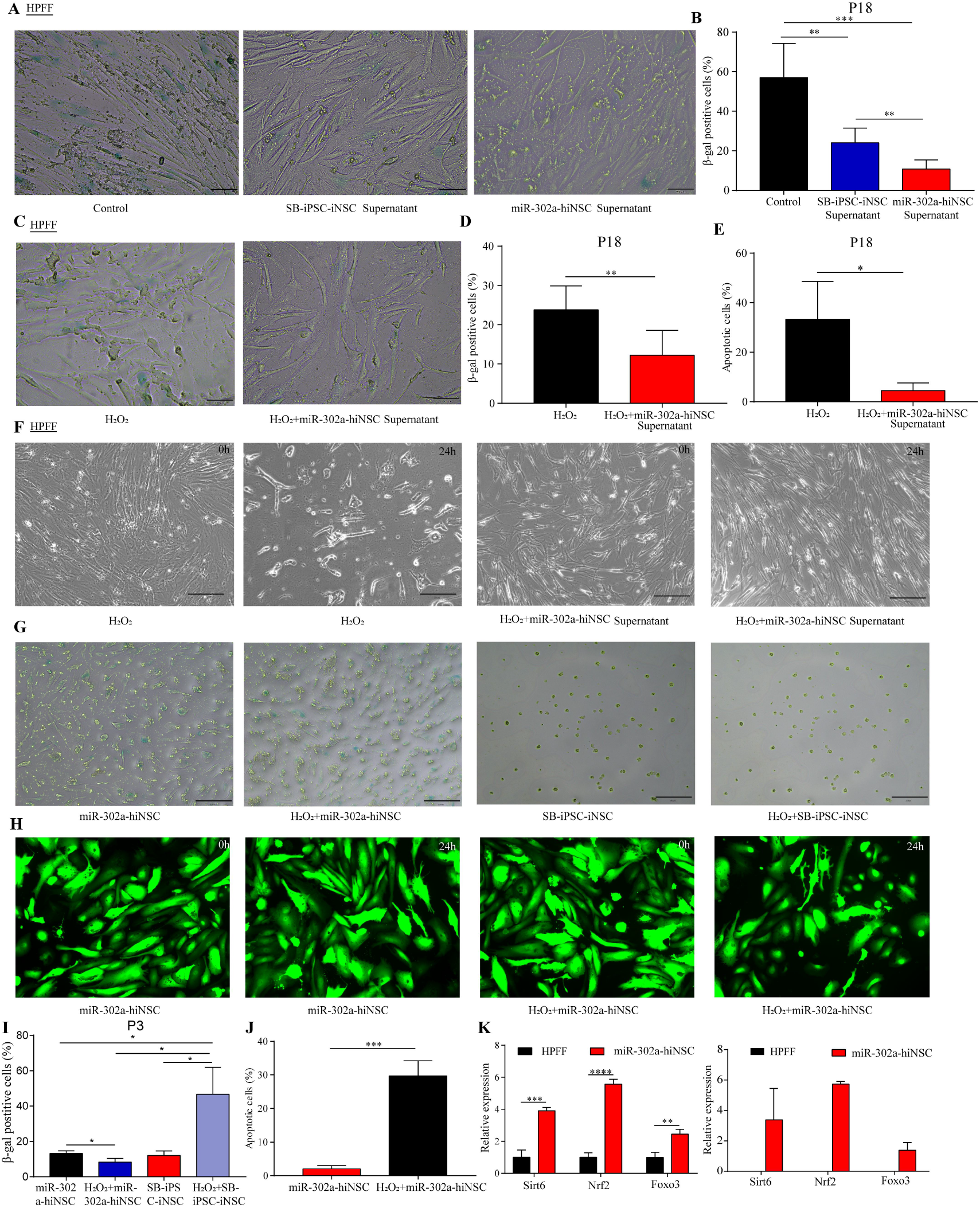
Anti-aging and antioxidant effects of culture supernatant from miR-302a-hiNSCs on senescent skin fibroblasts and neural stem cells. (**A**) β-gal staining analysis of HPFF, HPFF+SB-iPSC-iNSC Supernatant and HPFF+miR-302a-hiNSC Supernatant+H_2_O_2_ (P18). Scale bar, 100uM. (**B**) Quantitative analysis of β-gal staining of HPFF, HPFF+SB-iPSC-iNSC Supernatant and HPFF+miR-302a-hiNSC Supernatant+H_2_O_2_ (P18). Data are expressed as mean ± standard deviation. n=6, **p < 0.01, ***p < 0.001 (t-test). (**C**) β-gal staining analysis of HPFF+H_2_O_2_ and HPFF+miR-302a-hiNSC Supernatant+H_2_O_2_ (P18). Scale bar, 100uM. (**D**) Quantitative analysis of β-gal staining of HPFF+H_2_O_2_ and HPFF+miR-302a-hiNSC Supernatant+H_2_O_2_ (P18). Data are expressed as mean ± standard deviation. n=6, **p < 0.01 (t-test). (**E**) Quantitative analysis of apoptosis in HPFF treated with 300 mM H_2_O_2_ and HPFF cultured with miR-302a-hiNSC supernatant for 24 h. Data are expressed as mean ± standard deviation. n=6, *p < 0.05 (t-test). (**F**) Cell survival status under 24 h microscopy of HPFF treated with 300 mM H_2_O_2_ and HPFF cultured with miR-302a-hiNSC supernatant (P18). Specifically, HPFF cultured with miR-302a-hiNSC supernatant were resistant to H_2_O_2_-induced apoptosis. Scale bar, 200 uM. (**G**) Analysis of β-gal staining of miR-302a-hiNSC, SB-iPSC-iNSC, miR-302a-hiNSC +H_2_O_2_ and SB-iPSC-iNSC +H_2_O_2_ with 300 mM H_2_O_2_ for 24h (P3), scale bar, 200uM. (**H**) MiR-302a-hiNSC were treated with 300 mM H_2_O_2_, and the amount of GFP-positive cells was observed after 24 h. Scale bar, 100 uM. (**I**) Quantitative analysis of miR-302a-hiNSC, SB-iPSC-iNSC, miR-302a-hiNSC +H_2_O_2_, and SB-iPSC-iNSC +H_2_O_2_. Data are expressed as mean ± standard deviation. n=6, *p < 0.05. (**J**) Amount of GFP-positive cells after treatment of miR-302a-hiNSC with 300 mM H_2_O_2_ for 24 h. Data are expressed as mean ± standard deviation. n=3, ***p < 0.001 (t-test). (**K**) qRT-PCR analysis of the expression of three longevity factors Nrf2, Sirt6 and Foxo3 in HPFF, Ptf1a-hiNSC and miR-302a-hiNSC. y-axis indicates the relative mRNA expression of the three longevity factors. n=3, **p < 0.01, ***p < 0.001, ****p < 0.0001 (t-test).

### MiR-302a-iNSCs Improve Spatial Learning Ability and Memory in aged SAMP8 Mice

We tested whether transplanted miR-302a-hiNSCs had any therapeutic effect in the mouse SAMP8 model.After miR-302a-hiNSCs were implanted into the hippocampus of the SAMP8 mouse model, some behaviors were performed to evaluate the mice’s cognitive function at 4 and 8 weeks (14.5 and 15.5 months of age, respectively) (Fig. 6A). There were no differences in motor distance and motor speed between WT, SAMP8 and mice injected with miR-302a-hiNSCs in the 4w and 8w open-field experiments, while mice transplanted with miR-302a-hiNSCs spent less time at rest compared to WT and SAMP8 (Fig. 6, B and C). There was no difference in the number of times, time and total percentage of correct spontaneous alternate responses (SAP) to explore the new foreign arm in the Y-maze test (Fig. 6, F and G).

**Fig. 6.**
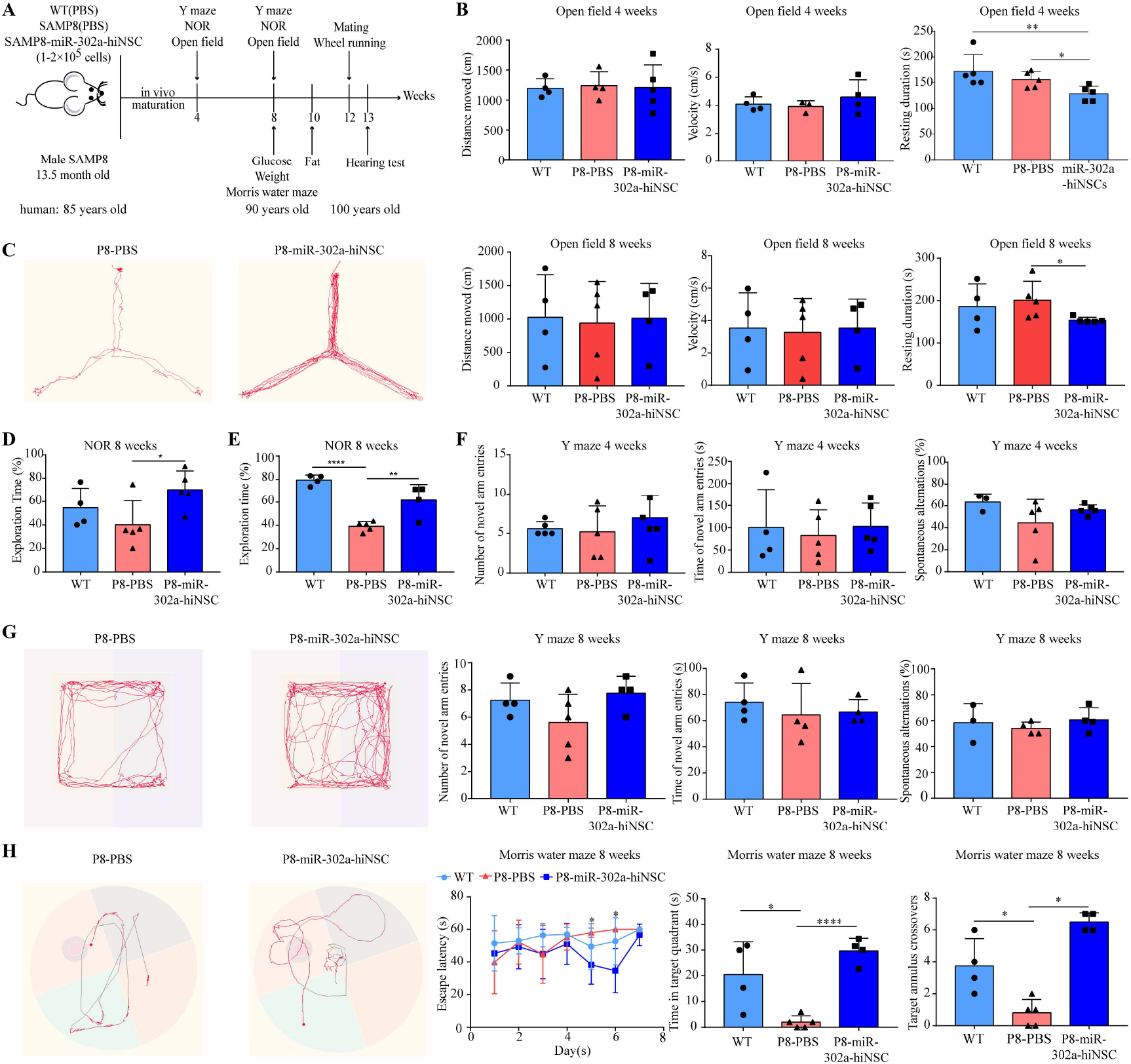
MiR-302a-iNSCs Improve Spatial Learning Ability and Memory in aged SAMP8 Mice. (**A**) Timeline for assessing the behavior of SAMP8 mice after transplantation of miR-302a-hiNSCs. (**B**) Open-field experiment fat 4 w. Results of the three-group mine field test in WT, SAMP8 and miR-302a-hiNSCs transplanted mice. *p < 0.05, **p < 0.01. (**C**) Open field experiment at 8w. Typical illustrative example of walking paths of SAMP8 and miR-302a-hiNSCs transplanted mice in the open field experiment in the mine field test. Results of three groups of WT, SAMP8 and miR-302a-hiNSCs transplanted mice in the mine field test. *p < 0.05. (**D**) New object recognition at 4 w. New object recognition test recorded as a percentage of time spent exploring new objects in the three groups. *p < 0.05. (**E**) New object recognition at 8w. The new object recognition test recorded the percentage of time spent exploring new objects for the three groups. **p < 0.01, ****p < 0.0001. (**F**) Y-maze at 4w. There was no improvement in the number, time and correct spontaneous alternating response (SAP) to explore the new dissimilar arm in the Y-maze. (**G**) Y-maze at 8w. Typical illustrative example of the walking pathway in SAMP8 and miR-302a-hiNSCs transplanted mice in the Y-maze test. number of times, time and correct spontaneous alternation response (SAP) to explore the new heterotopic arm did not improve in the Y-maze. (**H**) Morris water maze at 8w. Typical escape modalities in the Morris water maze test in SAMP8 and miR-302a-hiNSCs transplanted mice on day 7 hidden platform test. Learning curves for the Morris water maze acquisition test were obtained over a period of 6 days. 7d escape latency, time spent in the target quadrant and preference for target platform location were recorded for the Morris water maze experiment. *p < 0.05, ****p < 0.0001.

The WT group and mice injected with miR-302a-hiNSCs spent more time examining new objects during the new object recognition test than the SAMP8 group (Fig. 6, D and E), indicating that the SAMP8 mice injected with miR-302a-hiNSCs had a better short-term non-associative memory for the original environment.

We compared spatial learning and long-term memory between the three groups in the Morris water maze at 8 weeks post-transplantation (15.5 months of age). In all tests, we found significant differences in injected miR-302a-hiNSCs. SAMP8 mice showed latency deficits to find hidden platforms on days 5 and 6 of training compared to the WT group. At the same time point, mice injected with miR-302a-hiNSCs had a significantly lower mean escape latency than SAMP8 mice, indicating an improvement in spatial acquisition ability. Furthermore, mice injected with miR-302a-hiNSCs spent more time in the target quadrant and preferred the target platform location (Fig. 6H). These findings indicate that mice injected with miR-302a-hiNSCs have a strong memory for the previous location of the platform and improved spatial memory recovery. Finally, these findings suggest that injecting miR-302a-hiNSCs improves spatial learning and memory in the SAMP8 mouse model.

### MiR-302a-hiNSCs improves health span, fat, weight, blood sugar, hair, fertility and hearing in aged SAMP8 Mice

To summarize the lifespan, we estimated the survival rate. At 50% survival, the median survival was 12.7 months for untreated male SAMP8 mice and 17.5 months for miR-302a-hiNSCs-treated SAMP8 mice. The SAMP8-miR-302a-hiNSCs-treated male mice exhibited a 37.8% extension in median lifespan, equivalent to a mean survival of 102.5 years, with significantly longer survival (Fig. 7A).

**Fig. 7.**
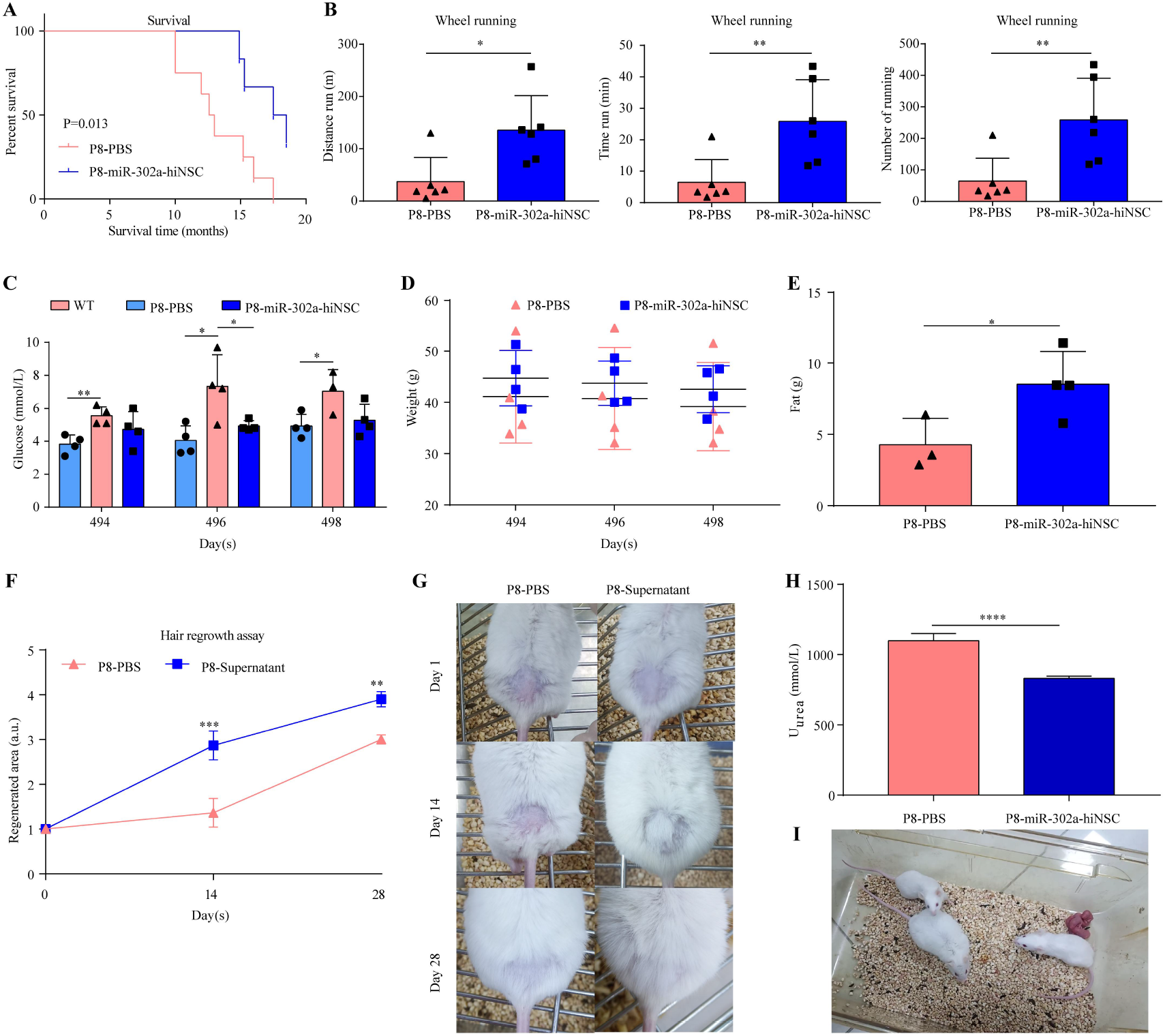
MiR-302a-hiNSCs improve fat, weight, blood sugar, hair and fertility in aged SAMP8 Mice. (**A**) Kaplan-Meier survival curves of SAMP8 mice and SAMP8+miR-302a-hiNSC mice (**B**) Wheel running experiment in 16.5-month-old mice. Distance, time, and number of running wheel runs. (**C**) Blood glucose levels in WT, SAMP8 and SAMP8+miR-302a-hiNSC mice fasted for 12h. (**D**) Quantification of body weight in SAMP8 and SAMP8+miR-302a-hiNSC mice. (**E**) Plot of fat content in SAMP8 and SAMP8+miR-302a-hiNSC mice. (**F**) Quantification of hair regeneration capacity of mouse dorsal skin. a.u., arbitrary units. (**G**) Typical images of hair regeneration in male SAMP8 mice before and after 28 days of treatment with supernatant of PBS/miR-302a-hiNSC. (**H**) Quantification of urinary urea changes in SAMP8 and SAMP8+miR-302a-hiNSC mice. (**I**) miR-302a-hiNSC improves fertility in SAMP8 mice. 16.5-month-old SAMP8+miR-302a-hiNSC mice are fertile after mating with BALB/c mice.

Physical activity is the main criterion of the frailty index in aged mice and is clinically associated with aging in humans. Therefore, in this study, the running wheel experiment was chosen as the physical activity parameter to be compared between the two groups. The results showed that 16.5-month-old male SAMP8 mice had decreased physical activity, compared with same-month-old SAMP8-miR-302a-hiNSCs male mice that exercised longer distances on the running wheel and spent more time and turned the wheel on the running wheel (Fig. 7B). Thus, miR-302a-hiNSCs significantly improved the healthy lifespan of mice, promoted physical activity performance, and reduced frailty in old age.

To further explore the normoglycemic effect of miR-302a-hiNSCs transplantation on the organism, the blood glucose of mice in the SAMP8 and SAMP8-miR-302a-hiNSCs groups was measured after 12 hours of fasting and repeated once every other day for a total of three times. The results showed that the blood glucose of the mice in the SAMP8 group was significantly higher, while the blood glucose of the mice in the SAMP8-miR-302a-hiNSCs group was obviously lower than that of the SAMP8 mice, which was not significantly different from that of the mice in the WT group at the age of 50 years and was within the normoglycemic range (Fig. 7C). Thus, miR-302a-hiNSCs could improve the blood glucose of SAMP8 mice and maintain a relatively young level.

Next, we measured the body weight and adiposity parameters of the two groups of mice. The SAMP8-miR-302a-hiNSCs group had a higher body weight but were not statistically different from the SAMP8 mouse group. The SAMP8-miR-302a-hiNSCs group was significantly different from the SAMP8 group in terms of adiposity (Fig. 7, D and E).

The ability of hair regeneration diminishes with age. To investigate the effect of miR-302a-hiNSCs on hair growth, we shaved approximately 1 cm × 1 cm square of hair on the back of each mouse on the day of transplantation. At day 14, we observed that the back hair regeneration area of the mice in the transplantation group recovered to about 80%, and at day 28, the back hairs of miR-302a-hiNSCs transplantation-treated SAMP8 mice had completely regenerated the entire area. In contrast, the hair regeneration capacity of PBS-injected SAMP8 mice was much lower (Fig. 7, F and G).

To evaluate the renal function of mice, we tested the urea concentration in the urine of two groups of mice at 17.5 months of age, and the urinary urea concentration of mice in the SAMP8-miR-302a-hiNSCs group was significantly lower compared to the SAMP8 group. Moreover, mice in the SAMP8-miR-302a-hiNSCs group could mate with females at 16.5 months of age (equivalent to 100 years of age in humans) and still had fertility (Fig. 7, H and I). To explore the effect of transplanting miR-302a-hiNSC on the auditory function of SAMP8 mice, we selected three test frequencies (8 kHz, 16 kHz, and 32 kHz) as representatives of the low, mid and high-frequency regions of the cochlea, respectively. At the 8 kHz test frequency, the hearing threshold was 60 dB in SAMP8 mice and 35 dB in SAMP8+miR-302a-hiNSC mice (Fig. S3A). At 16 kHz test frequency, the hearing threshold of SAMP8 mice was 55 dB and that of SAMP8+miR-302a-hiNSC mice was 25 dB (Fig. S3B). At 32 kHz test frequency, the hearing threshold of SAMP8 mice was 45 dB, and the hearing threshold of SAMP8+miR-302a-hiNSC mice was 25 dB (Fig. S3C). In conclusion, miR-302a-hiNSCs significantly prolonged the survival of SAMP8 mice, dramatically improved the healthy life span of mice, promoted physical activity performance, and improved the hair, fertility and hearing of SAMP8 mice.

## DISCUSSION

In this study, we demonstrate the use of a unique miR-302a reprogramming-based source of induced neural stem cells that promote cognitive improvement, life extension, and physical function in a murine model of late-stage severe AD. An Important feature of this cell is that it expresses Nrf2, Sirt6 and Foxo3 in addition to the fundamental neural stem cell signatures. Both human and murine fibroblasts can be employed to efficiently and rapidly establish iNSCs cells in about 2-3 days, and in vitro and in vivo studies have been performed to determine their function. These results support that autologous cell reprogramming into neural stem cell technology can be a viable approach to conventional autologous therapy without the necessity for invasive brain biopsies or lifelong immunosuppressive therapy.

We seek to utilize next-generation cell reprogramming to generate an easily isolated and autologous NSCs therapy to treat degenerative diseases. In a clinical setting, the potential for autologous cells to avoid immune rejection could provide additional therapeutic advantages over the allogeneic NSCs therapies currently used in clinical trials.

Human neural stem cells can differentiate into different types of neural cells, providing a potential source for repairing degenerated or damaged brains. However, human adult neural stem cells persist in the brain while encountering continuous challenges from age or injury-related changes in the brain environment (e.g., oxidative stress), gradually decreasing or depleting and becoming increasingly less functional. Aging is associated with cognitive decline and the development of multiple diseases such as Alzheimer’s disease, Parkinson’s, aging, and tumors (*9, 32, 33*). Therefore, the in vitro culture of human-induced neural stem/progenitor cells and their possible therapeutic applications have received particular attention (*27*).

In recent years, direct reprogramming techniques have been used to generate NPCs from murine and human somatic cells, which resemble neural stem cells in the brain (*34*). The reprogramming of iNSCs using techniques that involve pluripotent transcription factors such as SOX2 or OCT4 and non-neural pluripotent transcription factors such as BMi1, Zfp521, and Ptf1a faces issues including low induction efficiency and a lengthy process. (*21–26, 29, 35, 36*).

We use a single miR-302a reprogrammed fibroblast into direct iNSCs. In this single step, the rapid generation of large numbers of transplantable cells is critical for the cure of disease, as patients with severe degenerative disease or tumors often have little time left to live (*37*). In addition, the direct reprogramming process can eliminate the potential tumorigenicity of the iPS stage and help ensure that iNSCs do not form tumors in vivo. To further maximize the clinical translation potential of our study, we employed a single reprogramming strategy of miR-302a, which can substantially increase the generation rate of hiNSCs using microRNAs compared to existing reprogramming methods (*38*).

It is a new observation. Some studies have shown that miR-302 clusters can combine miR-367 to reprogram somatic cells into IPS with high efficiency (*38*). However, none of the studies showed that using a miR-302a alone can perform reprogramming somatic cells into neural stem cells. We determined that miR-302a alone can efficiently and rapidly reprogram fibroblasts into neural stem cells (hiNSCs) based on miR-302 cluster reprogramming. In vivo and in vitro cultures, hiNSCs can differentiate into both glial cells and neurons and does not express pluripotency markers. In vivo, hiNSCs survives for more than ffour weeks and remains positive for nestin or TUJ-1, but does not express OCT4. Moreover, this neural stem cell expresses Nrf2, Sirt6, and Foxo3, which have antioxidant and anti-aging properties, and the potential therapeutic use of the resulting human iNSC remains to be explored.

Research has shown that aging is the greatest risk factor for most diseases that affect people in later life, leading to a sustained decline in neural stem cells, cognitive decline, and the onset of some degenerative diseases, particularly Alzheimer’s disease with increasing age (*9*). As one of the top 10 deadly diseases worldwide, Alzheimer’s disease has a significant psychological, social and economic impact on patients, caregivers, families and society as a whole, and there is no cure for AD or a way to alter the course of the disease. Recently, stem cell transplantation has been proved to be an effective treatment for neurodegenerative diseases (*39, 40*).

The rate of decline in neurogenesis in the hippocampus appears to be accelerated in Alzheimer’s disease compared to physiological aging (*6, 41*). Similarly, in most mouse models of Alzheimer’s disease based on familial mutations, both SVZ and hippocampal neurogenesis are reduced, indicating a decrease in the number or impaired function of NSCs (*41, 42*). Although some effects of Alzheimer’s disease mutations can be reproduced in cultured NSCs, suggesting that Alzheimer’s disease mutations have a direct impact on NSCs function, other effects of mutations may also have indirect effects on NSCs through other cells. Indeed, risk factor genes involved in sporadic Alzheimer’s disease (e.g., APOE) impair astrocytes and microglia, which in turn can alter NSCs function (*43, 44*).

Many animal studies demonstrated that mesenchymal stem cells (MSCs), induced pluripotent stem cells (IPS), or NSCs have a positive role in AD treatment and may improve cognitive performance through a variety of immunological, histological, and genetic measures (*16, 27, 45–49*).

However, MSCs suffer from immune rejection and are not yet able to differentiate into corresponding neural cells and build neural connections at the damaged site, which affects their long-lasting effects. In contrast, autologous-induced neural stem cells do not have these problems and can also differentiate to form neurons. Functional integration of iNPC-derived neurons with host synaptic networks helps to enhance hippocampal plasticity and repair neural circuits in the host brain, thereby improving cognitive performance in AD mice (*27*). As NSCs repair capacity is limited in vivo, in vitro NSC supplementation can replenish NSC stocks to repair neurons lost after disease or injury.

Efforts to convert human somatic cells into induced neural stem cells (iNSCs) for cell replacement therapy have been extensive. In our study, we characterized the iNSCs generated through miR-302a and evaluated their effectiveness in an animal model of late-onset Alzheimer’s disease (AD). Studies have shown that AD is primarily a sporadic, late-onset disease with an exponential increase in prevalence starting at age 65. The SAMP8 mouse has several distinct advantages over genetically modified models and is particularly well suited to study the “transition” between aging and AD. The phenotype of the SAMP8 mouse resembles the symptoms of patients with late-onset and aging-related sporadic AD (*50*). Therefore, we used 13.5-month-old SAMP8 mice as an animal model of late AD for our study, an age equivalent to 85 years in humans. We subjected human iNSCs to hippocampal transplantation and showed significant improvements in dependent learning and memory abilities in three behavioral tests (minefield experiment, Y-maze, and Barnes water maze) at 4 and 8 weeks post-transplantation. This result suggests that human iNSCs generated based on miR-302a could be suitable for cellular therapy of AD (*50*).

Most diseases that impact the elderly population are primarily attributed to the aging process. Research suggests that various biological processes, including oxidative stress, telomere depletion, cellular depletion, and cellular senescence, play a role in age-related diseases (*51*). The decline in the number or deficit of NSCs significantly contributes to systemic aging (*13*). As a result, stem cell transplantation, exosome injection, anti-oxidative stress strategies, and telomere supplementation have been proposed to delay aging by supplementing or activating the relevant stem cells (*13, 17, 52*). Although stem cell transplantation can prolong life, the benefit is brief and requires the use of young stem cells (*13*). Exosomes must be injected multiple times, and the exosomes required are also derived from young stem cells. Older exosomes are useless, and exosome injections no longer work once the organism reaches a certain age (*17*).

Given this backdrop, the impact of miR-302a-derived iNSCs on senescence-affected tissues in rapidly aging mice remains to be explored. The present study aims to fill this knowledge gap by demonstrating that miR-302a-iNSCs have substantial anti-aging potential and that in vitro cellular experiments have shown that iNSCs to have more vital antioxidant and anti-aging effects than normal neural stem cells. Furthermore, in vivo experiments involving older SAMP8 mice revealed significant improvements in various aging-related functions following iNSCs injection, including increased lifespan, reduced frailty, hair regeneration, fat reduction, regulation of blood glucose, and restoration of renal and repair of reproductive function.

The median survival of male mice transplanted with iNSC’s SAMP8 was increased from 12.7 months to 17.5 months, representing a 34.8% life extension. Previous studies have shown that transplantation of neural stem cells can extend life by 10% in 8-month-old mice, equivalent to transplantation in humans at about 35 years of age. This experiment was implanted in SAMP8 mice transplanted at 13.5 months, equal to 85 years of age in humans, with a 34.8% life extension to a median survival of 17.5 months, which corresponds to an average survival of 102.5 years. The 34.8% life-extension result obtained with iNSCs was much higher than the 10% life-extension result obtained with anti-inflammatory neural stem cells. This result may be related to the expression of NRF2, Sirt6, and Foxo3 by these iNSCs. Because NRF2 has an antioxidant effect, it can promote the survival of neural stem cells and the life-prolonging effect of the organism (*53–56*). Engineered NRF2 can genetically enhance the modification of mesenchymal stem cells, and such GES cells are capable of dual resistance to cellular senescence and tumorigenic transformation (*57*). Sirt6 also has antioxidant effects, and overexpression of Sirt6 alone can extend the life span by 30% (*58*). Nrf2 and Sirt6 can be combined to promote the antioxidant effect of stem cells (*59*). Foxo3 also has an important role in stress resistance in neural stem cells (*32*). Moreover, Foxo3 genetically enhances vascular endothelial cells to fight aging and cancer (*60*).

Regarding physical strength index, the ageing SAMP8 mice transplanted with iNSCs were no worse than the young mice. Aging male SAMP8 mice transplanted with iNSCs ran farther on the running wheel, exercised longer, and were more resistant to fatigue than control SAMP8 mice.

We found that hiNSCs transplantation improved the deterioration of blood glucose due to aging and better-maintained serum glucose levels and energy homeostasis. This result is similar to that sirt6 overexpressing mice can restore blood glucose and energy homeostasis (*58*). The results suggest that Sirt6 expressed in iNSCs may be involved in glycemic regulation and energy homeostasis. It was found that iNSCs transplantation improved the increase in adiposity due to aging in SAMP8 mice and better maintained adipose homeostasis. This result also suggests that Sirt6 expressed in iNSCs may be involved in the regulation of lipid homeostasis. It is crucial to note that when we assessed renal performance based on urinary urea concentration, this could be a result of the decreased skeletal muscle catabolic rate and the change in urea content, indicating a considerable improvement in renal function. Additionally, we carried out mating studies on 16.5-month-old SAMP8, and we discovered that while no mice were born in the SAMP8-only group, four mice were born in the SAMP8 group transplanted with iNSCs. The outcomes showed that at the equivalent of 100 years in human life, the SAMP8 mice implanted with iNSCs were still capable of reproducing normally. Other experiments have not revealed anything similar.

In conclusion, our study highlights the potential of iNSCs generated based on miR-302a as a promising therapeutic approach for treating various age-related diseases and conditions. We found The iNSCs treatment to improve lifespan, cognitive abilities in late-stage Alzheimer’s disease, fatigue resistance, hair regeneration, blood glucose, and fat metabolism, renal function, reproductive function, and hearing loss. Further research will compare iNSCs from patient versus control donors, isolation and purification of iNSCs exosomes and optimisation of therapeutic pathways. These results pave the way for a novel approach to the rapid and efficient production of antioxidant and anti-aging hiNSCs for the preservation of cognitive function, improvement of aging, and prolongation of lifespan in Alzheimer’s disease patients.

## MATERIALS AND METHODS

### Human Samples

Fresh human foreskin samples from two healthy male donors (at the age of 26 and 29) were collected from Yuhuangding Hospital at Yantai, and fresh human eyelid sample from a healthy female donor (at the age of 41) was collected from Chang’ an Hospital at Xi*’* an. All the procedures were performed according to an IRB-approved protocol of either Binzhou Medical University or Northwestern Polytechnical University. The primary foreskin or eyelid cells were cut into pieces with ophthalmic scissors, enzymatically disassociated with (0.25% trypsin, 30~60 min), and cultured in fibroblasts culture medium, DMEM-High Glucose with 1% penicillin/streptomycin, 2 nM L-glutamine, and 10% FBS.

### Viral plasmid construction

The pGLV3-miR-302a plasmid was constructed using the lentiviral vector pGLV3/H1/GFP+Puro (pGLV3; Shanghai GenePharma Co., Ltd., Shanghai, China). The ADV3-U6-has-miR-302a-CMV-GFP overexpression adenoviral vectors plasmid also was constructed by Shanghai GenePharma Co., Ltd. The following sequence was used: miR-302a, 5’-TAAGTGCTTCCATGTTTTGGTGA-3’.

### Human and mouse iNSCs generation

Human fibroblasts isolated from the specimens were all expanded in the medium [DMEM/HIGH GLUCOSE+10% FBS+1% L-Glutamine+1% penicillin/streptomycin]. Human fibroblasts, derived from the foreskin or eyelid (at passages 3-10), were grown on glass coverslips coated with gelatin in a 24-well plate at a density of 1 x 10^4 cells per well in fibroblast culture medium. The following day, the medium was replaced with a fresh medium containing the miR-302a virus, polybrene (6 μl), and VPA (10 μg).Twenty-four hours post-transfection, Vitamin C (20 ug) was added within NSCs induction media 500 ul (DMEM/F12 + 2% B27 + 20 ng bFGF + 10 ng/mL EGF + 1% DOX + 1% L-glutamine+1% penicillin/streptomycin). After 72 h, Puromycin (10 uM) was added to remove untransfected cells, and then the media were changed into NSCs culture media (DMEM/F12 + 2% B27 + 20 ng/mL bFGF + 20 ng/mL EGF +1% L-glutamine+1% penicillin/streptomycin). The medium was changed every two days. From 24 h after transfection, some fibroblast morphology changed, and a few small colonies formed and expanded.

Mouse L929 cells were purchased from the ATCC and expanded in the MEF medium [DMEM/HIGH GLUCOSE (Hyclone) + 10% FBS (Gibco) + 1% L-Glutamine + 1% penicillin-streptomycin]. The procedure for the induction of mouse iNSCs (miNSCs) from L929 cells with miR-302a is essentially the same as for human iNSCs (hiNSCs). Mouse L929 cells were purchased from the ATCC and expanded in the MEF medium [DMEM/HIGH GLUCOSE (Hyclone) + 10% FBS (Gibco) + 1% L-Glutamine + 1% penicillin-streptomycin]. The procedure for the induction of mouse iNSCs (miNSCs) from L929 cells with miR-302a is essentially the same as for human iNSCs (hiNSCs).

### Real-time quantitative polymerase chain reaction (RT-qPCR)

Total RNA was isolated from iNSCs with the TaKaRa kit (Catalog# RR820A). RT-qPCR was performed with The Light cycle 96 Real-Time PCR System (Roche). The PCR-program: 95°C 30s, 95°C 5s, 60°C 30s. Steps 2-3 were repeated 40~45 times.

### Differentiation of hiNSCs and miNSCs

(1) Astrocytes differentiation: First, a 12-well plate was coated with 40 ug/mL poly-lysine for 1 hour at 37° C and washed with PBS 3 times. Then hiNSCs were plated onto those polylysine-coated wells, and astrocytes culture medium (DMEM/F12 medium supplemented with 10% FBS, 1 × NEAA, and two mM L-glutamine) was applied after hiNSCs attached to the bottom. (2) Oligodendrocytes differentiation: hiNSCs were enzymatically disrupted into single cells with trypsin in a 37 ° C incubator for 2 min and then resuspended and seeded onto a polylysine-coated glass coverslip and cultured in DMEM/F12 with 1 × N2, ten ng/ml PDGF, and three uM forskolin, half of the medium was changed every two days. Four days later, forskolin was substituted with 200 mM VC and cultured for seven days. (3) Neuron differentiation: For the generation of terminally differentiated neurons from hiNSCs, the cells were seeded at a density of 1 × 10^4^ cells per 12 mm glass coverslip coated with poly-ornithine/laminin in the neural induction medium (Neurobasal medium supplemented with 2% B27, 500 uM dbcAMP, 10 uM SB43152, 10 ng/mL BDNF, 10 ng/mL NT-3, 1 uM Ken, 2 mM Glutamax-ITM and 1% penicillin/streptomycin), half of the medium was changed every 2 days.

The procedure for differentiation of miNSCs is the same as for hiNSCs.

### Immunofluorescence staining

Cells were washed 3 times with PBS and then fixed with 4% paraformaldehyde (PFA, Sigma-Aldrich) for 10 min. Then cells were punched with 0.3% Triton for 10 min and blocked by 3% goat serum for 30 min at 37°C. After that, cells were incubated with primary antibodies overnight at 4°C followed by secondary antibody staining for 45 min at 37°C. After nuclear staining with Hoechst 33342 (Sigma-Aldrich), cells were observed and captured by Olympus CK51 microscope.

### Flow cytometry

INSCs were dissociated using trypsin. Cell suspensions were first fixed with 4 % formaldehyde for 10 mins on ice. After blocking for 30 min, cells were incubated with primary antibodies. After washing with PBS, cells were incubated with second antibodies. After another round of washing with PBS, FACS analysis was performed. Fluorochrome-matched isotype controls were used and subtracted during analysis.

### In vivo assay

Three weeks wild-type nude mice were anesthetized with 5% chloral hydrate at 10 uL/g. Fix the anesthetized mice on the brain stereotaxic apparatus and fix the microinjector needle, subtract the brain fur with scissors, cut the brain skin with a scalpel, expose the mouse head, wipe with hydrogen peroxide, adjust the microinjector needle to the fontanelle of the mouse brain with its zero point. 2 uL P3 hiNSCs cell suspension (100k) was microinjected into one side of the telencephalon (x: −0.22 cm, y: −0.26 cm, z: −0.14 cm) for 3 min using a micro-syringe. After injection, the needle was held for 3 min and then slowly and gently pulled out for 3 minutes. 2 to 4 weeks after transplantation, the frozen section of the brain was analyzed with immunofluorescence.

### MEA recording

For neuronal spontaneous action potential analysis on MEA plates, 24-well MEA plates containing 16 electrodes each were coated with 50 ug/mL laminin for three h at 37 ° C. The neurons derived from the differentiation of iNSCs were digested and centrifuged, the supernatant was removed, the amount of cell culture medium (neural induction media as previously described) added was adjusted based on cell density, and the proper culture medium was added and mixed. 30 μL of the cell suspension to each pore was added to ensure that all drops apply to the electrodes for 1 h at 37° C (25,000 cells in a wall), then added 250 uL of culture medium was to each well, and then 259 μL of culture medium, changing the media by half every three days. DIV 9 to 21-day-old MEA cultures were used for analysis. 5 min recordings/1h were performed every day except the day of media change using an MEA system (Axion Biosystems). We set the action potential spike-detecting threshold to 5.5 standard deviations.

### Cell Transplantation

In this study, 13.5-month-old male SAMP8 mice were used, purchased from the Animal Center of Peking University. GFP-labeled third-generation miR-302-hiNSCs were transplanted into the hippocampus’s dentate gyrus (DG) region in SAMP8 mice, with the following relative coordinates to the anterior fontanelle: AP, −1.06 mm; ML, ±1.0 mm; DV,-2.5 mm.

### Behavioral tests

Behavioral tests were performed on SAMP8, SAMP8 with transplanted cells, and age-matched WT mice. Before the behavioral tests, the mice were placed in the laboratory for 30 minutes to adapt. The open field and novel object recognition tests were performed before the Y maze and Morris water maze tests. Thus, the order of the behavioral tests was: open field test, novel object recognition test, Y maze test, Morris water maze test, and running wheel test.

### Open field test

The open field test setup consists of a white composite board measuring 40 cm in length, 40 cm in width, and 40 cm in height. The mice were placed in the center of the open field, each subject was placed in the same position, and a camera was fixed on top of the apparatus, recording the mouse’s movements for 5 minutes. And the total distance, speed, and immobility time were recorded automatically using Ethovision XT16. The testing apparatus was wiped with clean paper towels containing 75% ethanol between the two mice being tested.

### Novel object recognition test

On the first day, all mice were placed in the same equipment as the open field test to acclimate to the experimental environment in advance and were left alone to explore freely for 5 min. On the second day, two identical objects were placed in the apparatus, and each mouse was left to explore freely for 5 min. After 90 minutes, a new object was substituted for an old object, and the mice were allowed to explore for another 5 minutes, and the time spent exploring the old and new objects were recorded. The device was cleaned with 75% ethanol and wiped between the two subjects. The cognitive ability of the mice was tested by observing their exploration of old and new objects. The index observed was the new object recognition index, the time spent exploring new objects as T novel, exploring old objects as T old, and RI = T novel / (T novel + T old).

### Y maze test

The Y-maze comprises three arms, with an angle of 120° between each arm. The size of each arm (length x width x height) is 10 cm x 5 cm x 5 cm. First, cover the novel arm with a barrier, place the mouse in the Y-maze for 5 minutes, then remove the mouse and return it to the cage for 1 hour. After 1 hour, remove the barrier from the novel arm, place the mouse from the starting arm, and let it freely move in the three arms for 5 minutes. The spontaneous alternation rate is [(number of correct alternation responses) / (N-2)] × 100%.

### Morris water maze test

The Morris water maze experiment includes a circular water pool with a diameter of 110 cm and height of 40 cm, a camera system, a platform, and a system for analyzing animal behavior trajectories. Starting from the first quadrant of the water pool, if the mouse reaches the platform within 60 seconds, it is allowed to stay on the platform for 10 seconds. If the mouse does not find the platform within 60 seconds, it is placed on it and allowed to stay for 10 seconds. The same procedure is repeated for the other three quadrants, and the mouse is dried with clean tissue after completing the test. The platform is hidden for the next five days, and the mouse undergoes training just like the first day, but with the platform hidden underwater. After six consecutive days of pre-training, the platform is removed, and the testing of the mouse begins. During the testing, the mouse is placed into the water from the opposite quadrant of the platform’s original location and undergoes a 1-minute test. The video was recorded and analyzed using the Ethovision 16.0 software.

### Wheel running test

For the running wheel experiment, the mouse is first acclimated to the running wheel for three days at a speed of 8 rpm. On day 4, an incremental protocol is used: the running wheel speed starts at 8 rpm and increases to 10 rpm after 2 minutes. Fatigue is defined as the mouse’s unwillingness to move along the running wheel for at least 5 seconds despite mild electric shocks, and the distance and time of movement are recorded.

### Fasting blood glucose measurement in mice

The tail tip of the mouse is gently pricked with a blood glucose needle, and the fasting blood glucose is measured with a blood glucose meter. During the fasting period, the mouse is allowed to drink water freely.

### Body fat measurement

An instrument is used to measure the fat in the mouse to record the mass of lean tissue and fat.

### Urine Urea Measurement

The urine urea concentration in the mouse is calculated using a urea assay kit (Elabscience).

### Hair regrowth assay

On the first day, a 1 × 1 cm square of fur is shaved from the back of the mouse using a depilatory razor. Photos are taken after 14 and 28 days, and hair regrowth is evaluated using a 1-4 scale, with 4 representing complete hair regrowth, based on the photos and semi-quantitative assessment by two experimenters.

### Statistical information

In general, data for each result were obtained from 3 samples for cell culture and 8 animals for in vivo assay. Results are presented as the mean ± SD. Statistical analysis was performed using the Microsoft Excel computer programs and sigma 6.0. Unpaired two-tailed Student’s-tests and/or Mann-Whitney U-test were used to perform the difference in RT-qPCR results. P < 0.05 was considered statistically significant.

## Acknowledgments

We thank Ruonan Li for animal feeding, Renhui Zhan for RT-qPCR, Yanwei Wang for primary cell collection from human foreskin, Xuemei Hu for partial financial support and advice, Bo Dai for primary cell collection from human Eyelid and some writing advice, Yeying Sun for human IPS. The authors would like to thank Prof. Mengqing Xiang and Dr. Dongchang Xiao (Sun Yat-sen University) for providing the human and murine neural stem cells.

## Author contributions

Conceptualization: Y.S.-W. Methodology: Y.S.-W., Y.Y.-L., J.S., and Y.Y.-Z. Investigation: T.T.-X., and Y.N.-Z. Visualization: Y.Y.-L., J.S., and Y.Y.-Z. Supervision: Y.S.-W. Writing — original draft: Y.S.-W., and Y.Y.-Z. Writing — review & editing: Y.S.-W., and Y.Y.-Z.

## Data and materials availability

The data that support the findings of this study are available from the corresponding author upon reasonable request. Further information and requests for reagents may be directed to, and will be fulfilled by the lead contact Yuesi Wang.

## Supplementary Materials

Additional supporting information may be found online in the Supporting Information section at the end of this article.

