## Supplementary Materials for "MiR-302-Induced anti-aging neural stem cells enhance cognitive function and extend lifespan"

**This PDF file includes:**

Figs. S1 to S3
Tables S1 to S4

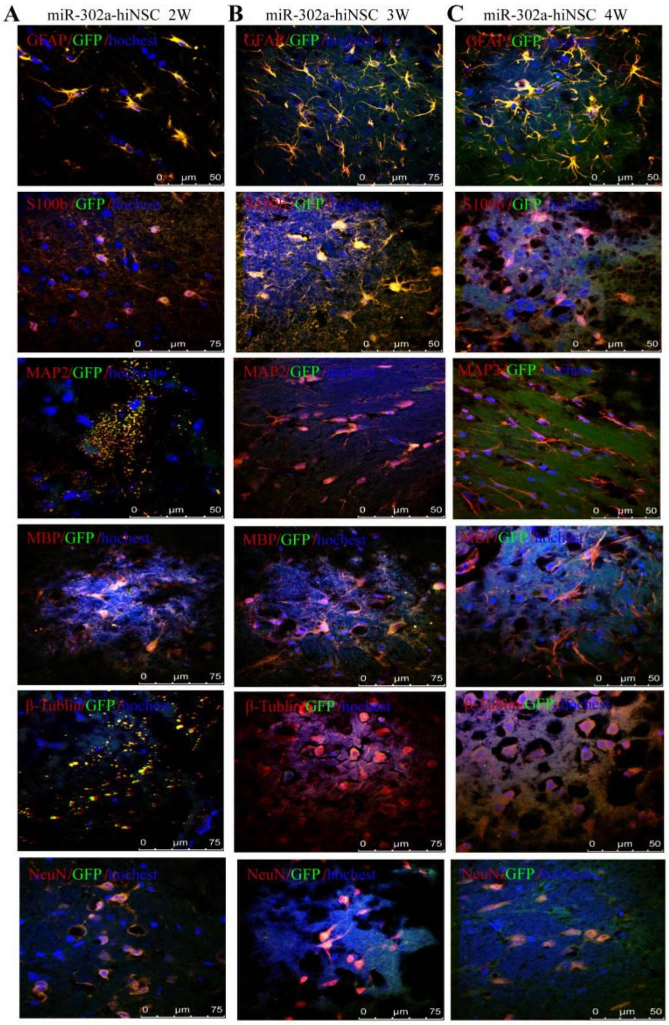


**Fig S1. Mulitipotency of miR-302a-reprogrammed hiNSCs in vivo.** (**A**) Immunostains reveal that hiNSCs can differentiate into GFAP+ and S100b+ astrocyte, Tuj1+ and NeuN+ neurons, and MBP+ and Olig2+ oligodendrocytes at 2 weeks after post-transplantation. (B) Immunostains reveal that hiNSCs can differentiate into astrocytes, neurons, and oligodendrocytes at 3 weeks after post-transplantation. (**C**) At 4 weeks after post-transplantation. GFAP+ and S100b+ astrocyte, Tuj1+ and NeuN+ neurons, and MBP+ and Olig2+ oligodendrocytes derived from hiNSCs could be detected by Immunostains.

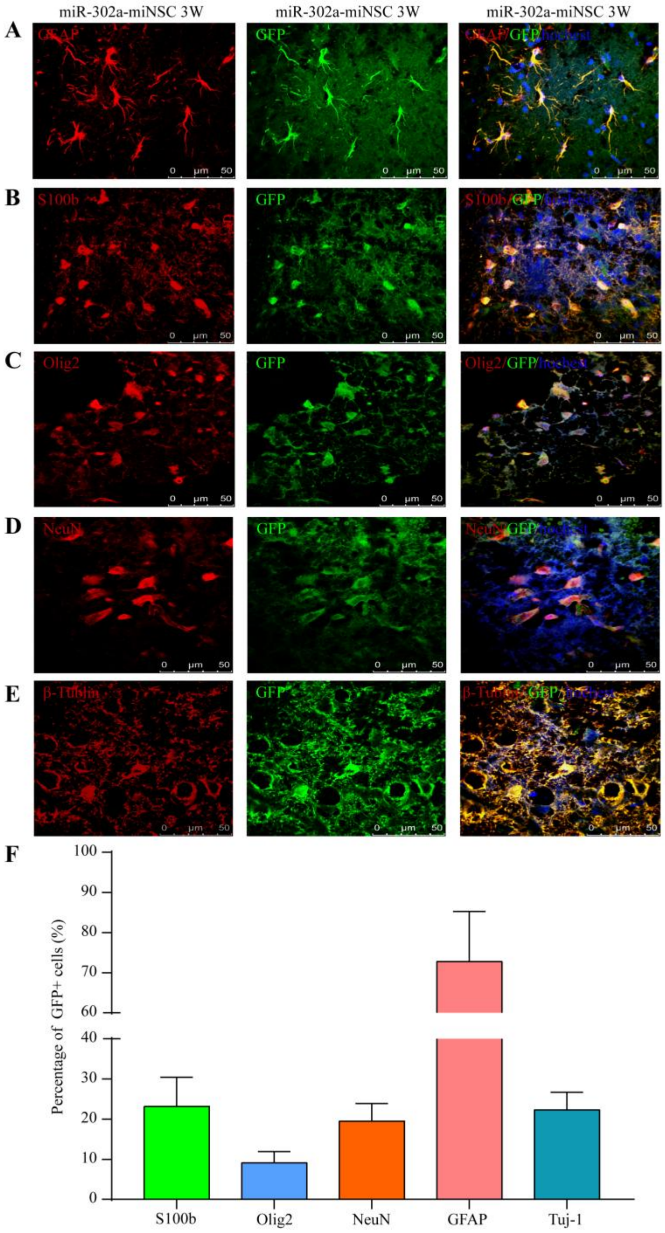


**Fig S2. Differentiation potential of miR-302a-reprogrammed miNSCs in vivo.** (**A** and **B**) Immunostains reveal that miNSCs can differentiate into GFAP+ and S100b+ astrocytes. (**C**) GFP+ miNSC can differentiate into Olig2+ oligodendrocytes. (**D** and **E**) Injected GFP+ cells have the neuron cell marker NeuN and Tuj1. (**F**) Quantification of labeled cells showed that GFP-positive cells differentiated into NeuN+ and Tuj-1+ neurons, GFAP+ and S100B+ astrocytes, and Olig2+ oligodendrocytes cells derived from miNSCs.

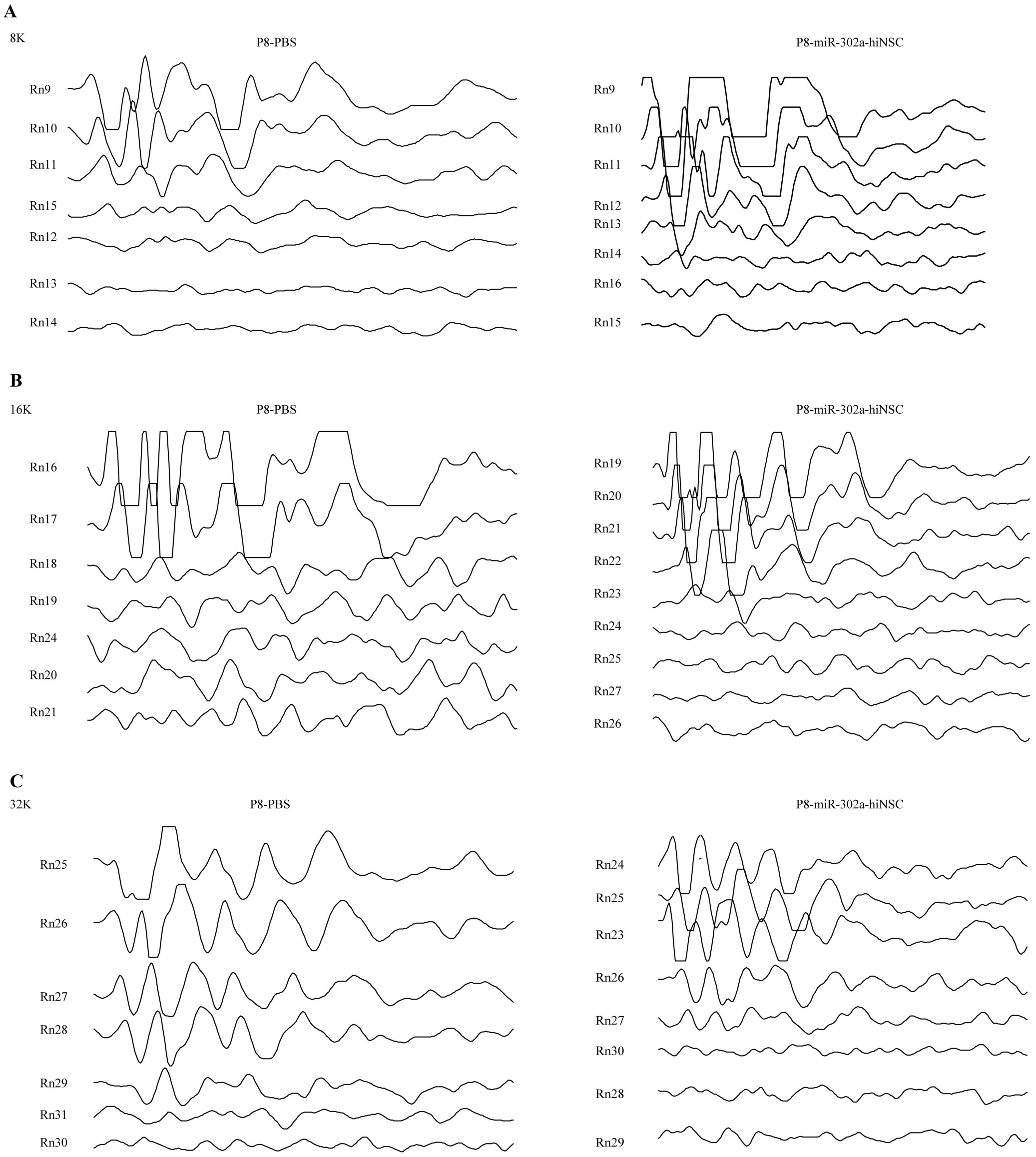


**Fig. S3. miR-302a-hiNSC improve hearing in aged SAMP8 mice.** (**A**) Hearing thresholds of SAMP8 and SAMP8+miR-302a-hiNSC mice at 8 KHz test frequency. (**B**) Hearing thresholds of SAMP8 and SAMP8+miR-302a-hiNSC mice at 16 KHz test frequency. (**C**) Hearing thresholds of SAMP8 and SAMP8+miR-302a-hiNSC mice at 32 KHz test frequency.

**Table S1. Overview of direct conversion methods for the derivation of human iNSCs.**

| Cell source chrom | Transcription Factor（s） | Others supplements / Pharmacological compound | Conversion efficiency | The earliest time of neurospheres emerge | Mean time to neurospheres formation | Clonal formation efficiency | Time of appearance of electrophysiological action potential | Reference |
|---|---|---|---|---|---|---|---|---|
| Human fibroblasts | miR-302a | | 86.8% | 13 hours | 2-3 day（d） | 3.04% （3 d）; 8.02%（7 d） | 10 day（d） | This submission |
| Mouse fibroblasts Human fibroblasts | SOX2 | | | 4 d | 6-10 d | | | Ring KL et al. cell stem cell,2012 |
| Human fibroblasts | SOX2 | | | 48 hours | 4 d | | | Bagó JR et al. Sci Transl Med, 2017 |
| human urine cells | OCT4, SOX2, SV40LT, KLF4 | miR-302/367 | | 12 d | 25 d | 0.2% | | Wang L et al. nature methods, 2013 |
| hDFs | SOX2 | let-7b，HMGA2 | | | 7-20 d | 0.2%-0.6% | | Kyung RY et al. Cell Reports , 2015 |
| Mouse fibroblast and human foreskin fibroblasts | Ptf1a | | | | | | 2 weeks | Xiao D et al. Nat. Commun,2018 |
| Cord Blood CD34+cells | OCT4 | | | | | | | Yang H et al. stem cells translational medicine,2015 |
| Adult human fibroblasts | OCT4 | | | 7+14 d | 3-4 weeks | | | Ryan R. M et al. Stem Cells,2014 |
| Human Neonatal and Adult Blood Cells | OCT4 | SMAD+GSK-3、(SB431542, LDN-193189, Noggin, CHIR99021) | | 8–10 d | | 0.024% | | Lee JH et al. Cell Rep, 2015 |
| human neonatal | | | | 24 d | 24-42 d | 0.4-0.7% | | Shahbazi E et al. |

| | | | | | | | | |
|---|---|---|---|---|---|---|---|---|
| (foreskin) fibroblasts (HNFs | Zfp521 | BAM groups | | | | | | Stem Cell Reports, 2016 |
| human UCB-MSC | SOX2 | | 25% | 14 d | | 0.015% | | Kim B et al. Cell Transplant,2018 |
| Human fibroblasts from AD patients，healthy person | SOX2 | 9 small molecules | 0.038%-0.091% | 12 d+6-8 d | 18-20 d | 0.09% | | Liu Y et al. J Alzheimers Dis, 2020 |
| Human Fibroblasts | OCT4 | RM | 0.94% | | | | | Ryan M et al. stem cells,2014 |
| adult human peripheral blood cells (PBCs) | SOX2，c-myc | CHIR99021、A83-01、hLIF、Trany1 | | 10 d | 10-21 d | 0.08% to 0.66% | 8–12 weeks | Sheng C et al.Nat. Commun,2018 |
| Human Dermal Fibroblast | c-MYC，SOX2 | | 0.2%–0.5 % | | | | | Daekee K et al. Mol Ther Nucleic Acids, 2019 |
| human neonatal dermis-derived fibroblasts or adult adipose-derived stem cells | OCT4, KLF4, SOX2, c-MYC | | 0.07%(neonatal HFF) 0.01%(adult HASC) | | 30-60 d | | 8 weeks | Dana M et al. Stem Cell Reports,2016 |
| human skin fibroblasts | OCT4, SOX2, Klf4, c-Myc | | | 17 d | | 0.20% | | Sandra M et al. Journal of Visualized Experiments,2015 |
| human fibroblast of amilial and Sporadic Parkinson's Disease Patients | OSKM | | | 7 d | 18-21 d | | | Lee M et al. Int J Stem Cells,2019 |
| Postnatal and adult human and monkey fibroblasts | OSKM | LIF, SB431542, CHIR99021 | 0.03%–0.08% | 13 d | 20 d | | 10 weeks | Lu J, et al. Cell Rep,2013 |
| Human PBMNCs | Klf4, OCT3/4, SOX2,c-MyC | | | 12 d | | | | Zheng W et al. J Vis Exp,2019 |
| Human fetal fibroblasts | SOX2，c-Myc | Brn2 or Brn4 | | 5-7 d | 6-7 d | 20-60 colonies / 10000 cells/well | 4–6 weeks | Qingjian Z, et al. The journal of biology chemistry,2014 |

| | | | | | | | | |
|---|---|---|---|---|---|---|---|---|
| Human bone marrow cells | Msi1，Ngn2，MBD2 | | | | | | | Vonderwalde I et al. Transl Stroke Res. 2020 |
| Human fibroblasts | SOX2, HMGA2, BRN4, SKM+SV40LT (BSKMLT) | phorbol-12-myristate-13-acetate, CHIR99021, SB431542 | | 7 d | 2 weeks | 1700 colonies/4 weeks | | Kwak TH et al. Int J Stem Cells,2020 |
| PBMC、ADFs、FPFs | BRN2, SOX2, KLF4, MYC, TLX, ZIC3 | BKSZ+CAPT | | 14 d | 19-24 d | 0.015-0.166% (19 d) | 10 weeks | Thier MC et al. Cell Stem Cell,2019 |
| human urine-derived cells | OCT4, SOX2, KLF4, GLIS1 | Purmorphamine, Forskolin, Vitamin C, Sodium butyrate (N) | | 8 d | 8-12 d | 11clonies/100, 000cells | 21 d | Kang PJ et al. Cells, 2019 |
| Human Fibroblasts | 25nTFs,15TFs, 13Tfs,6TFs,7TFs | | 10.54-11.22% | 6 d | | | 4-5 weeks | Pei SH et al. Stem Cell Reports,2017 |
| human fibroblasts | OCT4, or OCT4/SOX2/KLF4/p53shRNA | SB431542 and CHIR99021 | | | 2-3 weeks | | | Saiyong Z et al. nature protocols,2015 |
| Human bone marrow-derived cells, foreskin fibroblasts, keratinocytes | Msi1, Ngn2, and MBD2 | | 72 ± 8.97% and 86.25 ± 7.23% | 12 d | 2 weeks | | | Ahlfors JE et al. Stem Cell Res Ther. 2019 |
| adult human peripheral blood mononuclear cells | OCT4, SOX2, NANOG, LIN28, c-YC, KLF-4 and SV40LT | | | 10 d | 30 d | | 7 weeks | Xihe T et al. Stem Cell Research,2016 |

**Table S2. Overview of direct conversion methods for the derivation of mouse iNSCs.**

| Cell source | Transcription Factor（s） | Others supplements / Pharmacological compound | Conversion efficiency | The earliest time of neurospheres emerge | Mean time to neurospheres formation | Clonal formation efficiency | Time of appearance of electrophysiological action potential | Reference |
|---|---|---|---|---|---|---|---|---|
| Rat fibroblasts and astrocytes | Zfp521 or sox2 | | 46.1 ± 2.9% | | | | 6 weeks in vivo | Zarei-Kheirabadi M,et al .Stem Cell Res Ther,2019 |
| Mouse fibroblasts Human fibroblasts | SOX2 | | | 8 d | 6-10 d | 0.13%-0.96 % | 21 d | Ring KL et al. cell stem cell,2012 |
| Mouse fibroblast | Ptf1a | | | 6 d | 9-14 d | 0.5% at 14 day | 3 weeks | Xiao D et al. Nat. Commun,2018 |
| Mouse astrocytes | ZFP521 | | 41.7 ± 6.2% | | | | 60 d | Zarei-Kheirabadi M, et al. J Cell Physiol. 2019 |
| MEFs | Sox2, Klf4, c-Myc, Oct4 | | | 11 d | 19 d | 11 neurosphere /130,000 cells | 3 weeks | Thier M, et al. Cell Stem Cell,2012 |
| MEF | Oct4, Sox2, Klf4, c-Myc | | 0.07% | 11 d | 11-15 d | 0.69%-0.5% | 20 d | Kim J, et al. PNAS,2011 |
| mouse fibroblasts | Sox2, Klf4, c-Myc, Brn4 | | | 4-5 week | | | | Kim SM, et al. Nature protocols,2014 |
| mouse fibroblasts | BSKM. Brn4, Sox2, Klf4, c-Myc | | 6.3 ± 0.43% | 4 weeks | 4-5 weeks | | 14-16 d | Kim SM, et al. J. Biol. |

| | | | | | | | |
|---|---|---|---|---|---|---|---|
| | (BSKM) | | | | | | Chem,2016 |
| mouse fibroblasts | Brn4,Sox2,Klf4,c-Myc, plus E47/Tcf3 | | 4-5weeks | 4-5weeks | 1-5 clusters/5× $10^4$ cells | 7-16 d | Han DW, et al. cell stem cell,2012 |
| C57BL/6MEFs; C3H MEFs | Brn4,Sox2, Klf4,c-Myc (BSKM) | 2.99% (C57BL/6 MEFs) 8.22%(C3H MEFs) | | 6 week | | | Kim SM, et al. Stem Cell Research,2016 |
| mouse fibroblasts | Ezh2,Jarid2,Mtf2, Nanog,Pou5f1,Sall4,Smarca4, Sox2, Suz12, Tcf3 | | | | | | Yaqubi M, et al. Stem Cell Research & Therapy,201 |
| mouse fibroblasts, Liver Cells,B Lymphocytes | Brn2,Hes1,Hes3,Klf4,Myc,Notch1,(NICD),PLAGL1, Rfx4 13TFs；14Tfs | 1.00% | | 30 d | | 4 weeks | Cassady JP, et al. Stem Cell Reports, 2014 |
| mouse fibroblasts | Oct4,Sox2,Klf4,c-myc,Brn2,FoxG1,et al.11 factor | 12.3 % | 24 d | 25 d（13d FoxG1+ Sox2） | 3-317 colonies/20000 cells | 25 d | Lujan E, et al. PNAS, 2011 |
| Astrocyte’s mouse | SOX2 ASCL1 | 23.2%± 5.3% | | | | | Niu W, et al. Stem Cell reports,2015 |
| sertoli cells | Sox2,Pax6, Ngn2, Hes1， Id1, Ascl1, Brn2.c-Myc, and Klf4 | 0.70% | | | 0.0002% | 3 weeks | Chao S, et al. Cell Research,2012 |

**Table S3. Gene specific PCR primers used for RT-qPCR analyses.**

| Primer | | Primer sequences (5'-3') |
|---|---|---|
| GAPDH | Forward | GGAGCGAGATCCCTCCAAAAT |
| GAPDH | Reverse | GGCTGTTGTCATACTTCTCATGG |
| Nestin | Forward | CTGGAGCAGGAGAAACAGG |
| Nestin | Reverse | TGGGAGCAAAGATCCAAGAC |
| OCT4 | Forward | CCTCACTTCACTGCACTGTA |
| OCT4 | Reverse | CAGGTTTTCTTTCCCTAGCT |
| SOX2 | Forward | CCCAGCAGACTTCACATGT |
| SOX2 | Reverse | CCTCCCATTTCCCTCGTTTT |
| PAX6 | Forward | ATGTGTGAGTAAAATTCTGGGCA |
| PAX6 | Reverse | GCTTACAACTTCTGGAGTCGCTA |

**Table S4. Antibodies for immunofluorescence staining.**

| Antibodies | Company | Catalog# | Dilution |
|---|---|---|---|
| Nestin | BOSTER | BA1289 | 1:300 |
| OCT4 | Bioss | Bs-0830R | 1:300 |
| PAX6 | Bioss | Bs-11204R | 1:300 |
| SOX2 | Bioss | Bs-0523R | 1:300 |
| GFAP | BOSTER | ZA-0117 | 1:300 |
| Olig2 | BOSTER | ZA-0561 | 1:300 |
| β-Tub | ZSBIO | ZM-0439 | 1:300 |
| MBP | BOSTER | BA0094 | 1:300 |
| MAP2 | BOSTER | BM1243 | 1:300 |
| MAP2 | Bioss | Bs-20265R | 1:300 |
| Synapsin 1 | BOSTER | BA1421-2 | 1:300 |
| VIM | BOSTER | PB0378 | 1:300 |
| CY3 | BOSTER | BA1031 | 1:200 |
| IgG | BOSTER | BA1038 | 1:200 |
| Dylight549 | Abbkine | A23320 | 1:200 |
| Dylight549 | Abbkine | A23310 | 1:200 |
